# Data-driven simulations elucidate how lymphocyte motility behaviors influence cell-cell interactions within germinal centers

**DOI:** 10.1101/2025.08.05.668700

**Authors:** Nikita Sivakumar, Chanhong Min, Kibaek Choe, Dhruv Seth, Chris Xu, Wendy Béguelin, Jude M. Phillip, Feilim Mac Gabhann

## Abstract

Within the germinal center light zone, B cells and T cells rely on cell motility to engage in B-T interactions that enable antibody affinity maturation. Prior studies have shown that disrupting baseline lymphocyte motility behaviors can impair the germinal center response. However, the quantitative relationship between lymphocyte motility and B-T interactions in the germinal center is not well understood. A key challenge is that many *in vivo* imaging approaches require sparse cell labeling to segment and track individual cells within the dense germinal center microenvironment. This sparse labeling approach obscures a substantial fraction of B-T interactions within the tissue and limits the ability to map measured cell motility behaviors to emergent cell-cell interactions. A second challenge is that cell motility is intrinsically heterogeneous, with cell populations exhibiting a range of behaviors that are difficult to systematically relate to multi-cell interactions. To address these limitations, we developed a computational framework that allows us to 1) quantify heterogenous motility behaviors from intravital imaging data and 2) systematically predict how these measured behaviors lead to emergent cell-cell interactions. Applying unsupervised clustering, we identified seven distinct lymphocyte motility behaviors (B1-B7) from intravital imaging data. Notably, T-cells tended to take on fast, exploratory movements (B1-B4) while B cells tended to move in slower, confined paths (B5-B7). We then developed and validated a three-dimensional agent-based model to quantify the effect of these individual motility behaviors on B-T interactions. Our model predicted that B cells exhibiting slow motility behaviors (B5-B7) and T cells exhibiting fast motility behaviors (B1-B4) maximized the emergent number of unique B cell interactions per T cell within fixed observation volumes. These predictions suggest that the baseline differential motility behaviors of B cells and T cells enhance their ability to surveil for B-T interactions during the germinal center response.

## INTRODUCTION

Lymphocyte motility drives cell-cell interactions that enable key elements of the adaptive immune response.^1–3^ Disruption of baseline lymphocyte motility behaviors can lead to defective cell-cell interactions,^4,5^ and ultimately dysregulate the adaptive immune response. Systematically understanding how distinct motility behaviors drive dynamic cell-cell interactions would yield critical insights into how researchers could tune motility properties to modulate critical immune cell-cell interactions. Here, we present a computational pipeline that leverages intravital imaging to learn *in vivo* lymphocyte motility behaviors and agent-based simulations to predict how these motility behaviors impact multicellular interactions.

To develop this pipeline, we focused on how B cell and T cell motility drive emergent cell-cell interactions during the peak of the germinal center reaction. Germinal centers (GCs) are polarized, multi-cell structures that form in secondary lymphoid organs to enable production of high affinity antibodies in response to antigen exposure.^6–11^ Within the dark zone compartment of GCs, germinal center B cells (GCBs) undergo rapid proliferation and somatic hypermutation to generate distinct antibody clones.^6^ Dark zone GCBs then migrate to the GC light zone, where GCBs use surface antibodies to internalize antigen presented by follicular dendritic cells (FDCs). GCBs then present this antigen to T follicular helper cells (Tfhs) with cognate T cell receptors, initiating a GCB-Tfh interaction that provides critical cues for clonal selection and antibody affinity maturation (e.g. selection cues determine whether the GCB will mature into an effector cell, re-enter the dark zone for further rounds of hypermutation, or undergo apoptosis).^9,10^ Light zone, GCB-Tfh interactions are a key mechanism of antibody selection during the adaptive immune response. GCBs and Tfhs rely on cell motility to navigate the GC light zone and engage in these GCB-Tfh interactions.^12–15^ Prior studies have shown that dysregulated lymphocyte motility in bulk cell populations leads to diminished antibody responses.^4,5^ However, the relationship between lymphocyte motility patterns and GCB-Tfh interactions within the GC light zone is not well understood. Here, we leveraged intravital imaging of mouse GCs along with a validated computational pipeline to systematically quantify this relationship.

Multi-photon intravital time-lapse microscopy enables live-cell imaging of lymphocyte motility and interactions within *in vivo* germinal centers.^16–19^ This imaging approach captures high-resolution movies of single-cell GCB and Tfh trajectories by selectively labelling a subset of GCBs and Tfhs within the mouse GC, thereby allowing these cells to be distinguished from the dense GC microenvironment. This sparse labeling approach yields high-quality data of single-cell trajectories and interactions between labelled cells. However, there are two key limitations to intravital imaging studies: 1) sparse labelling obscures interactions between labelled and unlabeled cells (which often comprise the majority of the imaged volume) and, therefore, measured GCB-Tfh interactions may be skewed by sampling biases; and 2) because intravital imaging only captures a small sub-volume of the imaged tissue at high resolution, measured trajectories are skewed towards those that start and end within this imaging volume.^20^ Therefore, while we can gather insights about lymphocyte motility and GCB-Tfh interactions from intravital imaging, sparse labeling introduces sampling biases that prevent systematically mapping measured motility behaviors to interactions using these data alone.

In this study, we developed and validated an agent-based model (ABM) that integrates insights from intravital imaging data to predict how single-cell lymphocyte behaviors drive emergent GCB-Tfh interactions. Agent-based models simulate how several, individual cell agents following stochastic rules can drive emergent phenomena at the population level.^21–24^ This bottom-up structure enables ABMs to link dynamic single-cell behaviors, such as cell motility or cell adhesion, to multi-cell outcomes, such as cell-cell interactions or cell spatial patterning.^25–29^ While prior studies have leveraged ABMs to understand how GCB and Tfh cell behaviors contribute to the GC response,^30–33^ these studies modelled lymphocyte movement as a random walk with the same bulk speed and turning properties. Our group has shown that cell motility is inherently heterogeneous and bulk measurements across populations of cells can overlook the distinct motility patterns taken on by a single cell type.^34,35^ Here, we aimed to expand prior work in this field by 1) developing a parameterization approach that allows our ABM to simulate the single-cell heterogeneity of observed lymphocyte motility patterns and 2) applying the ABM to predict how these distinct motility patterns impact emergent, multi-cell interactions. By constructing our ABM to accurately recapitulate the observed heterogeneity of lymphocyte motility behaviors within GCs, we can systematically explore our ABM to map single-cell motility properties to emergent B-T interactions.

To accomplish these goals, we developed **PRISMM** (**<u>P</u>**ipeline for **<u>R</u>**ecapitulating Cell-Cell **I**nteractions using **<u>S</u>**patial **<u>M</u>**otility **<u>M</u>**odeling, **Figure 1**). PRISMM first applies unsupervised clustering to identify emergent motility behaviors across single-cell, GCB and Tfh trajectories measured from intravital imaging. Unlike bulk analysis approaches, this single-cell approach assesses the heterogeneity of movement patterns that a cell may exploit to perform key biological functions.^34^ PRISMM then uses a three-dimensional ABM to simulate these learned cell motility behaviors in multi-cell environments and predict how they influence the likelihood of GCB-Tfh interactions. Critically, PRISMM uses a parameterization approach that enables the ABM to model the heterogeneous properties of each individual motility behavior quantified from the experimental data. We validated the ability of PRISMM to accurately recapitulate each observed motility behavior and to predict emergent GCB-Tfh interactions. Using this pipeline, we reconstructed the movements and interactions of cells at the full, expected GC density, and predicted the ‘unseen’ interactions between unlabeled cells. We also predicted how a finite imaging volume can undercount the abundance of fast-moving cells and tested the extent to which this sampling bias impacts predicted interactions. Finally, we systematically explored how individual observed GCB and Tfh motility behaviors influenced GCB-Tfh interactions. Our computational pipeline directly leverages insights from intravital microscopy data to uncover how cell motility drives GCB-Tfh interactions (**Figure 1**).

**Figure 1.**
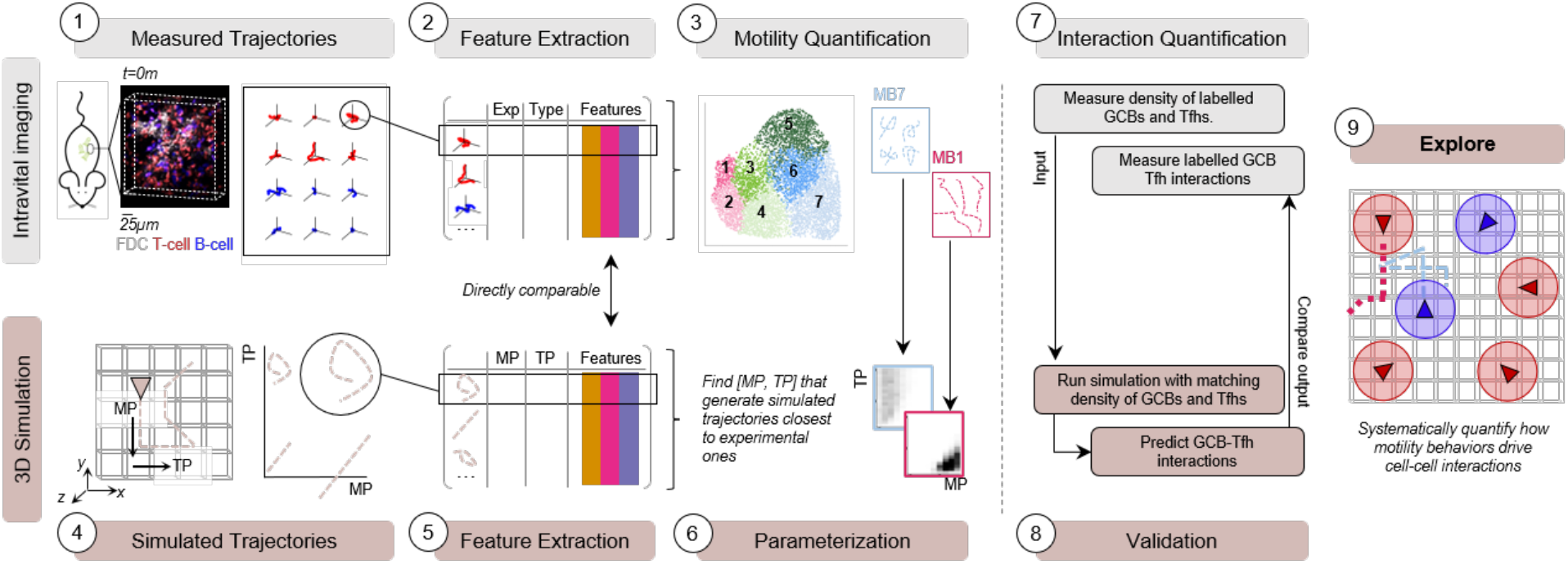
Schematic of PRISMM pipeline with parallel analysis of experimental and computational data. **(1)** PRISMM first extracts single-cell GCB and Tfh trajectories from time-lapse, intravital imaging of *in vivo* mouse germinal centers. **(2)** From each lymphocyte trajectory, PRISMM calculates several motility features (Figure 2, **Table 1**). **(3)** PRISMM then applies unsupervised clustering on the multi-dimensional motility feature space to identify distinct motility behaviors (MBs). **(4)** In parallel, PRISMM uses a 3D, on-lattice agent-based model (ABM) to simulate cell motility based on parameters that control speed and tortuosity (move-probability (MP) and turn-probability (TP), respectively). This parameter space can be uniformly sampled to generate a universe of simulated trajectories. Because the model is stochastic, we simulate many trajectories per simulation parameter set. **(5)** We calculate the same motility features as in (2) from each simulated trajectory so that the simulated trajectories are directly comparable with experimental trajectories. **(6)** PRISMM then fits (MP, TP) parameter distributions to recapitulate each experimentally measured motility behavior. To validate the ABM, **(7)** we ran simulations that recapitulated the labelled densities of GCBs and Tfhs from each experimental video, and **(8)** tested whether the simulation-predicted number of GCB-Tfh interactions, using the experimental cell densities as input, was comparable with measured interactions in each video. **(9)** Finally, we applied the mechanistic simulations to systematically explore how distinct motility behaviors drive GCB-Tfh interactions.

**Table 1.**
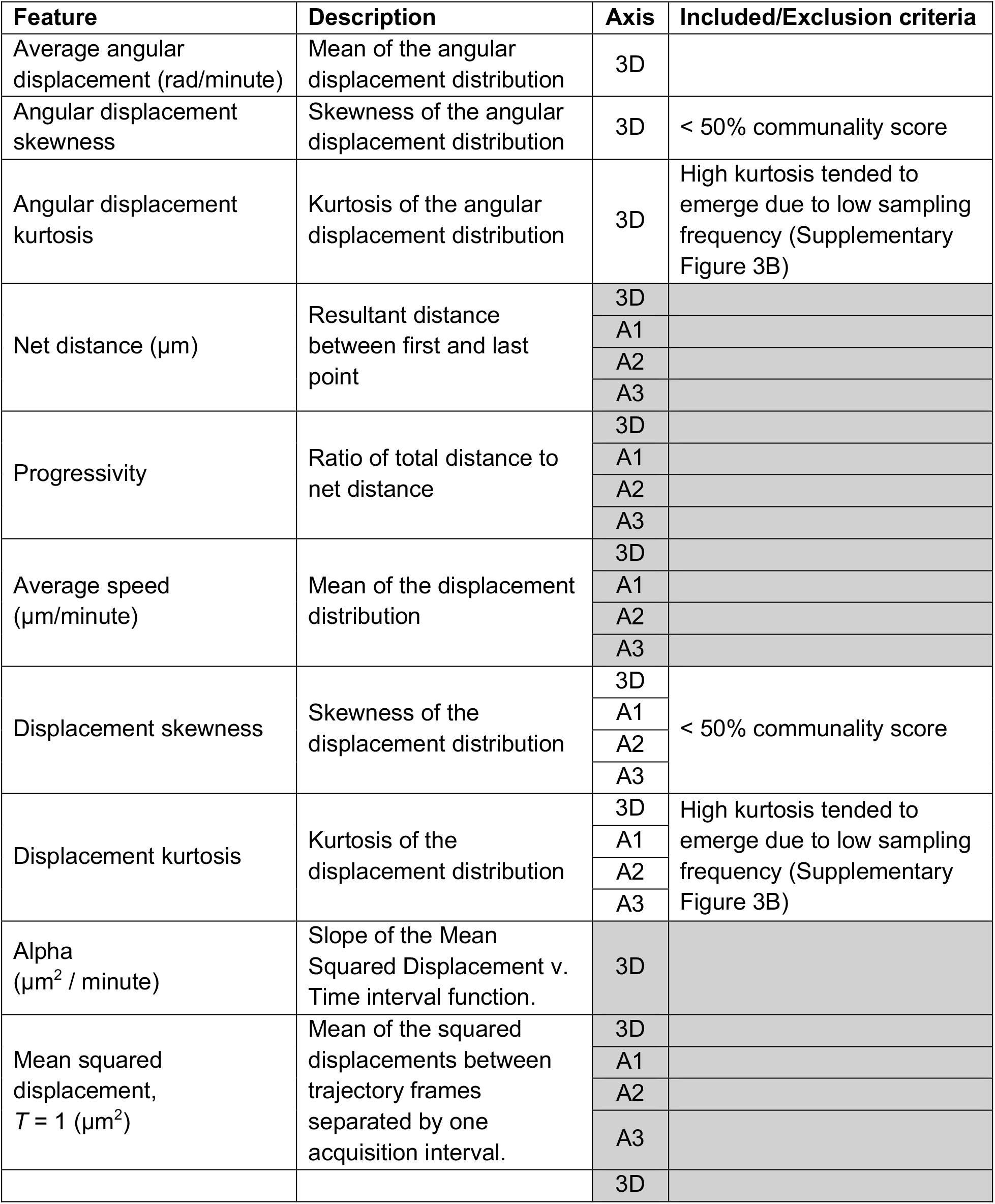
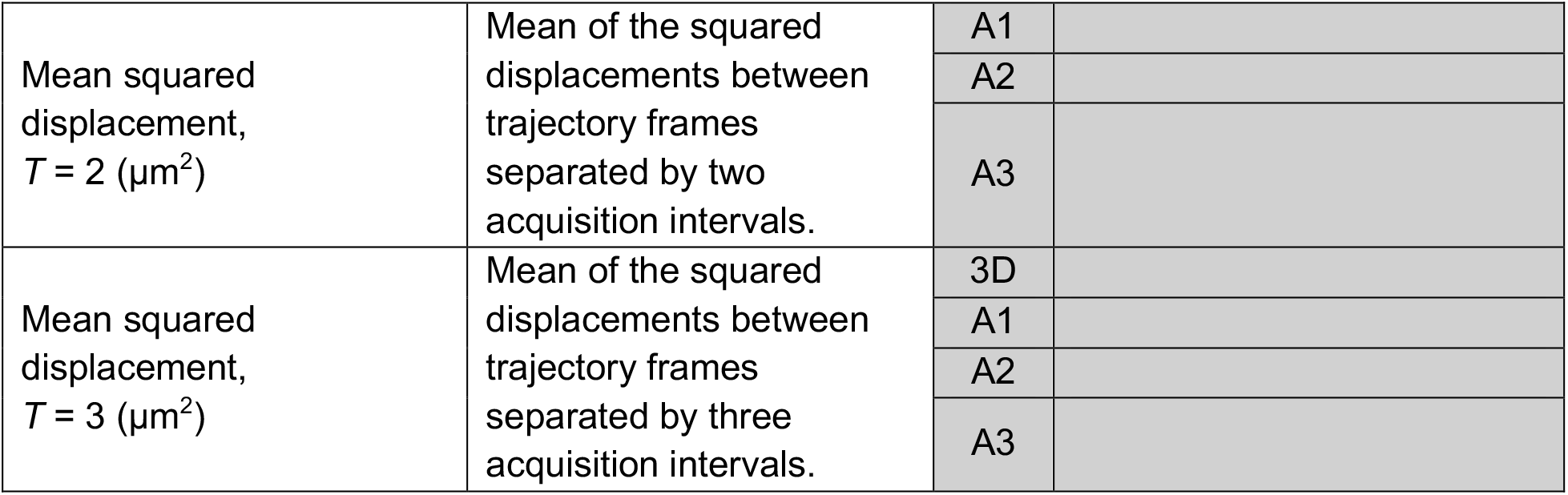
Motility features computed for experimental and simulated trajectories. Each described feature was computed for each trajectory. Included features are highlighted in gray. Criteria for excluded features are described.

| Feature | Description | Axis | Included/Exclusion criteria |
|---|---|---|---|
| Average angular displacement (rad/minute) | Mean of the angular displacement distribution | 3D |  |
| Angular displacement skewness | Skewness of the angular displacement distribution | 3D | < 50% communality score |
| Angular displacement kurtosis | Kurtosis of the angular displacement distribution | 3D | High kurtosis tended to emerge due to low sampling frequency (Supplementary Figure 3B) |
| Net distance (µm) | Resultant distance between first and last point | 3D |  |
|  |  | A1 |  |
|  |  | A2 |  |
|  |  | A3 |  |
| Progressivity | Ratio of total distance to net distance | 3D |  |
|  |  | A1 |  |
|  |  | A2 |  |
|  |  | A3 |  |
| Average speed (µm/minute) | Mean of the displacement distribution | 3D |  |
|  |  | A1 |  |
|  |  | A2 |  |
|  |  | A3 |  |
| Displacement skewness | Skewness of the displacement distribution | 3D | < 50% communality score |
|  |  | A1 |  |
|  |  | A2 |  |
|  |  | A3 |  |
| Displacement kurtosis | Kurtosis of the displacement distribution | 3D | High kurtosis tended to emerge due to low sampling frequency (Supplementary Figure 3B) |
|  |  | A1 |  |
|  |  | A2 |  |
|  |  | A3 |  |
| Alpha<br>(µm<sup>2</sup> / minute) | Slope of the Mean Squared Displacement v. Time interval function. | 3D |  |
| Mean squared displacement,<br>*T* = 1 (µm<sup>2</sup>) | Mean of the squared displacements between trajectory frames separated by one acquisition interval. | 3D |  |
|  |  | A1 |  |
|  |  | A2 |  |
|  |  | A3 |  |
|  |  | 3D |  |
| Mean squared displacement,<br>*T* = 2 (µm<sup>2</sup>) | Mean of the squared displacements between trajectory frames separated by two acquisition intervals. | A1 |  |
|  |  | A2 |  |
|  |  | A3 |  |
| Mean squared displacement,<br>*T* = 3 (µm<sup>2</sup>) | Mean of the squared displacements between trajectory frames separated by three acquisition intervals. | 3D |  |
|  |  | A1 |  |
|  |  | A2 |  |
|  |  | A3 |  |

## RESULTS

### Intravital imaging enables single-cell tracking of GCBs and Tfhs within the germinal center light zone

To analyze the motility behaviors of germinal center lymphocytes, we performed two-photon (2P) intravital microscopy on bone marrow chimeric mice with fluorescently labelled Tfhs and GCBs. We used the Cγ1-cre strain to drive GC-specific expression of YFP and CFP reporters in GCBs, and CD4-cre to drive the tdTomato reporter in Tfhs. We imaged mouse popliteal lymph nodes at the peak of the germinal center reaction (9-11 days post-immunization with NP(4-hydroxy-3-nitrophenylacetate) ovalbumin) to acquire single-cell motility data and cell-cell interaction data between Tfhs and GCBs (**Supplemental Table 1**). We captured three-dimensional (3D) time-lapse images (231 µm x 231 µm x 90 µm volume) at 30-second intervals. Imaging duration ranged between 80-90 minutes (161 - 181 frames). We then used IMARIS to segment and track individual cells within each time-lapse movie. To maximize the number of tracked cells, we tracked cells for 9.5 minutes (20 frames).

### Tfhs and GCBs exhibit distinct motility phenotypes

To characterize GCB and Tfh motility, we applied a single-cell approach that allows us to identify 1) whether the lymphocytes exploit distinct motility patterns and 2) the extent to which each cell type exploits distinct combinations of these patterns.

To accomplish this task, we quantified several motility features from each 3D trajectory to characterize the distributions of displacement and angular displacement for each lymphocyte (**Figure 2**, **Table 1**). After data quality control (see *Methods***, Supplemental Text S1** and **Supplementary Figures 1-3**), we performed a factor analysis to select features that explain critical variance across the observed trajectories (**Supplementary Figure 4A**). Factor analysis selected 26 motility features that included average speeds and mean-squared displacements (MSDs) in 3D and along the primary, secondary, and tertiary axes; net distance in 3D and along the primary axes; and progressivity in 3D and along the primary axes (**Supplementary Figure 4B**, **Table 1**). Excluded features comprised the skewness values of the displacement distribution, which described negligible variance across the observed trajectories. We constructed a two-dimensional (2D) UMAP embedding to visualize the 26-dimensional motility feature space of the 6,912 observed trajectories. To determine whether Tfhs and GCBs had distinct motility behaviors, we colored each trajectory in the UMAP by cell type. We found that Tfhs and GCBs occupied distinct subspaces within the 2D UMAP (**Figure 3A**). Displacement features appeared to horizontally differentiate trajectories in the UMAP space (**Figure 3B**), while tortuosity features appeared to vertically differentiate trajectories in the UMAP space (**Figure 3C**). Tfhs and GCBs were horizontally separated in the UMAP space (**Figure 3A**). Accordingly, Tfhs had significantly higher average speeds and MSDs at one-minute intervals than GCBs (**Figure 3B**). Tfhs and GCBs were distributed across a wide vertical range in the UMAP, suggesting that both cell types exhibited a broad range of tortuosity characteristics. Tfhs exhibited significantly lower progressivity than GCBs and significantly higher net displacement than GCBs (**Figure 3C**). This trend suggests that, on average, Tfhs are more motile and travel longer distances than GCBs.

**Figure 2.**
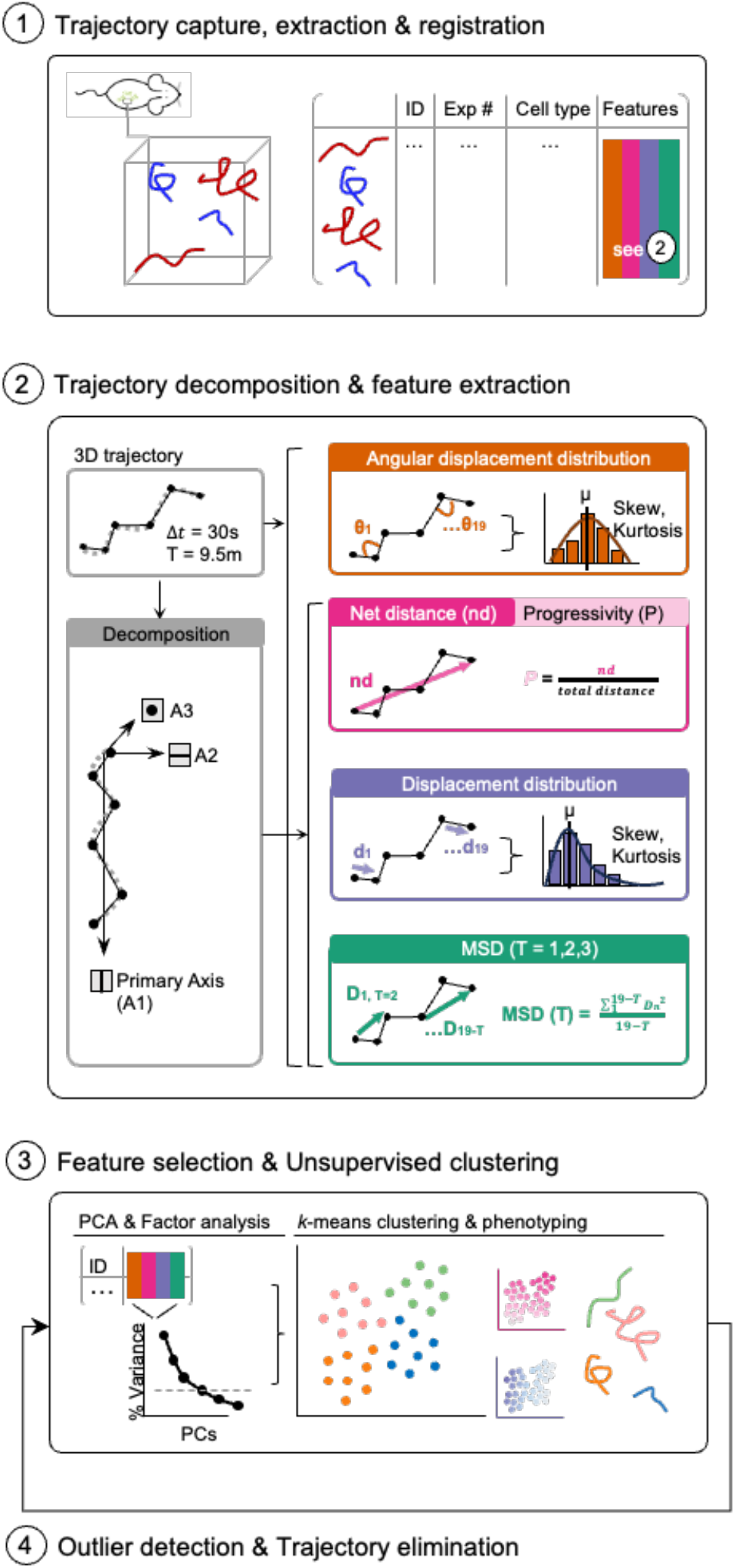
Schematic of quantified motility features for each trajectory. **(1)** From the intravital microscopy data, we segmented and tracked 3D trajectories for individual GCBs and Tfhs. Each trajectory comprised the (x,y,z) coordinates of a single cell in 20 consecutive frames acquired at 30-second intervals. **(2)** For each experimental trajectory, we decomposed the trajectory into primary, secondary, and tertiary axes of movement. For the 3D trajectory, we quantified statistical features of the angular distribution. For the 3D trajectory and the trajectory decomposition, we quantified the following features: net distance and progressivity; statistical features of the displacement distribution; and the mean squared displacement at intervals of 1, 2, and 3 frames (30 seconds, 1 minute, and 1.5 minutes, respectively). **(3)** After extracting these features, we projected the multi-dimensional feature space into a UMAP embedding and applied unsupervised clustering to identify motility behaviors. We then examined the behavior of each motility cluster. **(4)** Finally, we eliminated possible outlier trajectories and repeated step (3) on the cleaned dataset.

**Figure 3.**
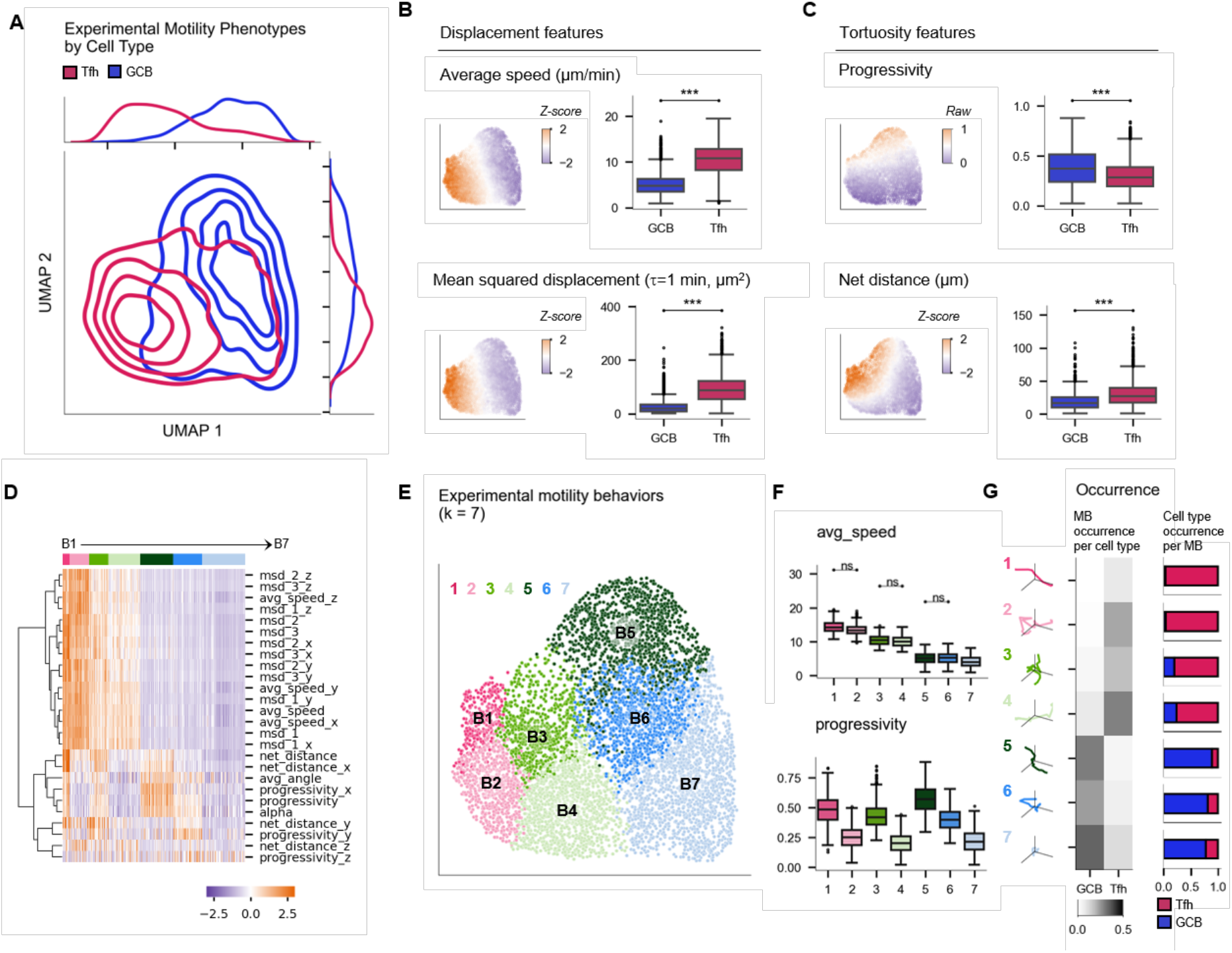
Quantification of distinct GCB and Tfh motility behaviors. **A,** Two-dimensional UMAP embedding of the motility features for GCB and Tfh trajectories (26 motility features per 6,912 trajectories). Lines on contour plots indicate 20% intervals of the cell populations. Embedding represents 3,184 Tfhs and 3,728 GCBs. All other UMAP projections shown in the figure correspond to this UMAP. **B,** Projection of standard-scaled average speed (top, left) and mean squared displacement at 1-minute intervals (bottom, left) onto UMAP embedding. Corresponding boxplots of average speed (top, right) and mean-squared displacement (bottom, right) for each cell type. **C,** Projection of raw progressivity values (top, left) and standard-scaled net distance (bottom, left) onto UMAP embedding. Corresponding boxplots of progressivity (top, right) and net distance (bottom, right) distribution for each cell type. **D,** heatmap representing standard-scaled values of each motility feature (rows) for each experimentally measured trajectory (columns). Rows are hierarchically clustered using Euclidean distance and Ward linkage. Trajectories are ordered by *k-means* cluster label. The *k*-means cluster label is shown on the top color band. **E,** Projection of seven motility behaviors (B1-B7) derived from *k*-means clustering onto the UMAP embedding. **F,** Boxplots of average speed and progressivity value distribution for each motility behavior. All comparisons were statistically significant (*p*<0.05) except for those marked by “n.s.” **G,** Representative trajectories for each motility behavior (left-most). Proportion of each motility behavior within each cell type (left heatmap) and proportion of each cell type within each motility behavior (right bar plot). Statistical testing shown on boxplots with multiple groups indicates results from a Kruskal-Wallis test, followed by a Dunn test for post-hoc comparisons. Statistical testing shown on boxplots with two groups indicates results from a Mann-Whitney test. * indicates *p*<0.05, ** indicates *p*<0.01, and *** indicates *p*<0.001.

To group the trajectories into distinct motility behaviors, we performed unsupervised *k*-means clustering of the motility feature space.^34^ This method divided the observed trajectories into seven motility behaviors (B1-B7, **Figure 3D-E**). We used inertia and concordance analyses to optimize the number of clusters that minimize within-cluster variance without overfitting (see *Methods* and **Supplementary Figure 4C-D**). Each motility behavior had characteristic displacement and tortuosity distributions, suggesting that the measured lymphocytes exploit distinct movement behaviors with varying speed and turning characteristics (**Figure 3F, Supplementary Figure 4E**). Tfhs and GCBs exhibited distinct combinations of these motility behaviors. Tfhs had a higher occurrence across B1-B4 (fast-moving clusters), while GCBs had a higher occurrence across B5-B7 (slow-moving clusters) (**Figure 3G**). Taken together, this result shows that Tfhs and GCBs exhibit distinct motility behaviors with different combinations of movement and turning.

### Calibration of 3D agent-based model to mimic simulate single-cell motility behaviors that mimic observed lymphocyte properties

After identifying distinct motility behaviors across GCBs and Tfhs, we wanted to quantify how these distinct motility behaviors influence GCB-Tfh interactions. Because unlabelled GCB-Tfh interactions are not observed within the intravital data, we cannot elucidate this relationship using the imaging data alone. Therefore, we developed and validated an agent-based model to simulate the observed motility behaviors within a multi-cell environment and systematically predict how single-cell motility properties influence multi-cell interaction dynamics.

We developed a 3D, on-lattice ABM in which cell agents execute movement rules at each timestep. Since displacement and tortuosity features were key differentiators of the experimental trajectories, we designed the cell movement rule with two parameters that can directly modulate these motility features. On each timestep, the cell agent evaluates a move probability (MP), the likelihood of moving forward, and a turn probability (TP), the likelihood of turning. When a cell agent turns, that cell agent will choose to turn to one of the 26 neighboring voxels, at random, that is not in the direction to which the cell is currently pointing. Each cell agent has its own MP and TP values. Because this movement rule is probabilistic, an (MP, TP) parameter combination can yield one of several possible trajectories. To use the ABM to recapitulate experimentally measured trajectories, we explored how simulated parameters affect distinct motility features, calibrated the simulation spatiotemporal resolution, and then optimized specific (MP, TP) parameter distributions to recapitulate each experimentally observed motility behavior (**Figure 4A**).

**Figure 4.**
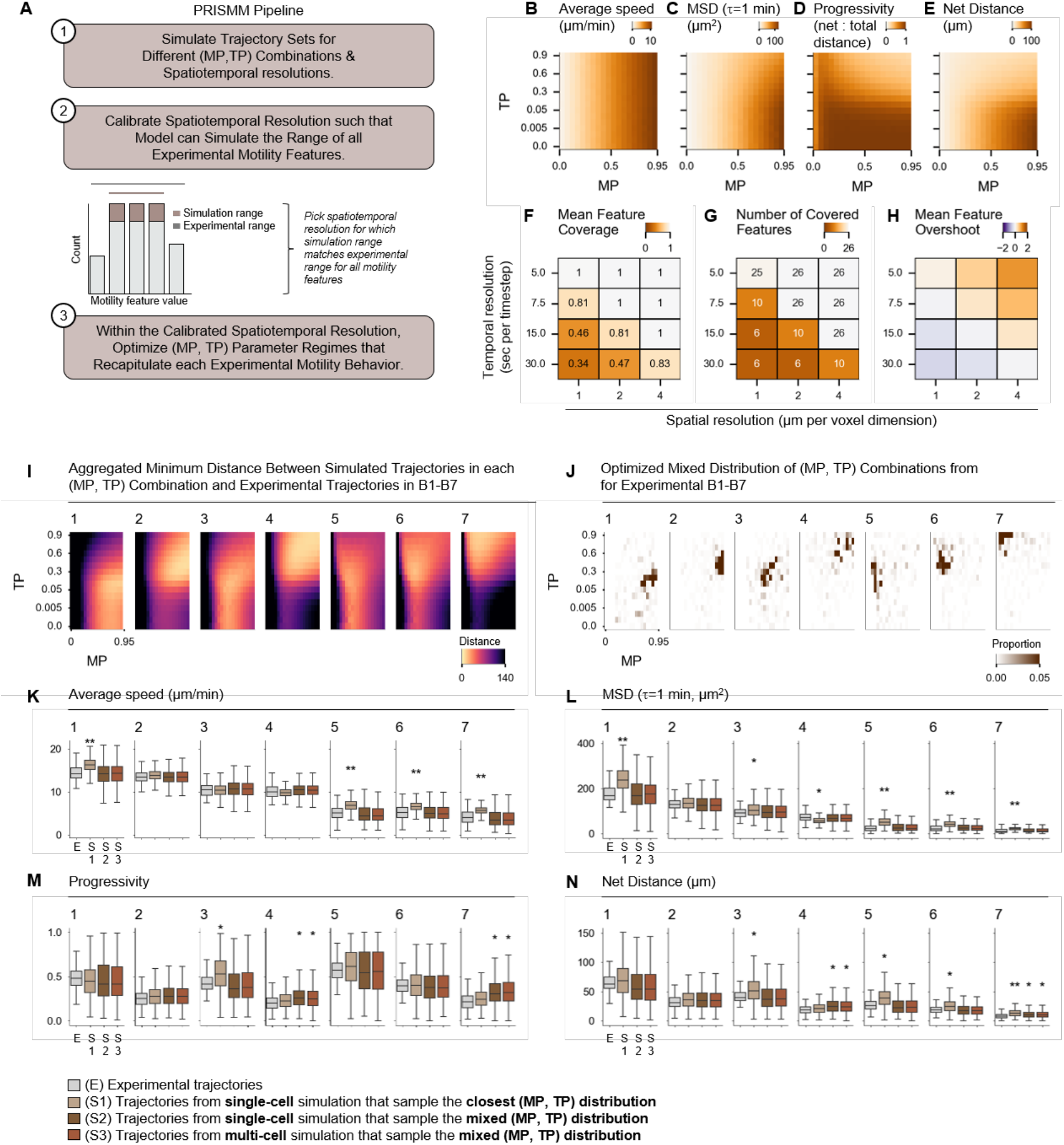
Calibrating ABM to recapitulate experimentally measured motility behaviors. **A,** Flowchart describing the process for developing the ABM that PRISMM employs to mechanistically simulate experimentally measured motility behaviors. **B-E:** Heatmaps representing average values across 2000 trajectories generated per (MP, TP) combination in single-cell simulations for the following motility features: average speed **(B)**, mean squared displacement at 1-minute intervals **(C)**, progressivity **(D)**, and net distance **(E)**. **F,** Heatmaps representing the average proportion of the experimental range captured by the simulated range across 26 features, at various spatial and temporal resolutions. **G,** The number of motility features (out of 26 total) for which the simulated range encompassed the experimental range, at different spatial and temporal resolutions. **H,** The log_2_ of the average ratio between the simulated range and experimental range across all 26 features for each spatial and temporal resolution. **I,** Heatmaps representing the aggregated minimum distance between the 2,482 simulated trajectories for each (MP, TP) parameter set in the multi-cell ABM and the experimentally measured trajectories within each experimental motility behavior. **J,** Heatmaps representing the optimized mixed proportions of (MP, TP) parameter sets to recapitulate each experimentally measured motility behavior. **K-N,** Boxplots comparing motility feature distributions for each motility behavior between the experimentally measured trajectories (“E”), simulated trajectories in the closest (MP, TP) parameter set (“S1”), simulated trajectories in the optimized mixed distribution of (MP, TP) parameter sets (“S2”), and simulated trajectories from multi-cell simulations resampling this parameterization (“S3”) for the following motility features: average speed **(K)**, mean squared displacement at 1-minute intervals **(L)**, progressivity **(M)**, and net distance **(N)**. * indicates that the distribution has a Cohen’s D value > 0.5 when compared with the experimental distribution. Multi-cell simulations represented in this figure were run with periodic boundary conditions for a total volume of 55 x 55 x 55 voxels, a spatial resolution of 2µm per voxel, and temporal resolution of 7.5 seconds per timestep.

To quantify how MP and TP affected the motility features of simulated trajectories, we ran single-cell simulations for distinct MP and TP values. Single-cell simulations model individual cell agent movement without additional rules to prevent overlap with other cell agents (see Methods). Because these single-cell simulations are more computationally efficient than multi-cell simulations, we initially used this simplified model to explore how model parameters impact single-cell motility behaviors. We sampled MP and TP from 0 to 1 in simulations with a voxel size of 4 µm and a temporal resolution of 30 seconds per timestep. The average speed (**Figure 4B**) and MSD at one-minute intervals (**Figure 4C**) increased directly with MP, such that higher MP values corresponded to greater simulated displacement. The progressivity of simulated trajectories decreased with respect to TP, such that higher TP values generated more tortuous trajectories (**Figure 4D**). The net distance depended on both MP and TP; the net distance of the trajectory increased as MP increased, while the net distance decreased as TP increased (**Figure 4E**).

In addition to MP and TP, the simulated spatial and temporal resolution of the ABM also affect cell agent movement. Our on-lattice simulation assumes that cells can only move between adjacent lattice spaces. The spatial resolution indicates the size of each voxel in the simulation lattice environment and controls how far a cell can move on a discrete timestep of the simulation. The temporal resolution indicates how long each simulation timestep represents and, thus, controls the number of opportunities a cell agent has to move (i.e., higher temporal resolutions necessitate longer run times to match the experimental trajectory duration, resulting in a greater possible number of moves). As the spatial resolution increases, the range of simulated displacements decreases. Conversely, as the temporal resolution increases the range of simulated displacements increases (**Supplemental Text S2** and **Supplementary Figure 5**). Prior to optimizing (MP, TP) combinations that can recapitulate observed experimental motility behaviors, we had to calibrate the spatiotemporal resolution of our ABM such that the possible ranges of motility features that could be generated from our ABM match that of those observed in the experimental data.

To calibrate the spatiotemporal resolution of the ABM, we compared the range of each motility feature for the aggregated simulated trajectories from a given spatiotemporal resolution to that of the experimental trajectories. We computed three metrics to assess the extent to which simulated motility feature distributions encompass the ranges of experimentally measured motility feature distributions (**Figure 4F-4H**). The mean feature coverage indicates, on average, the fraction of the experimentally measured range that the simulation set captures for each motility feature; ideally, we want this metric to be as close to 1 as possible (**Figure 4F**). The number of covered features indicates the number of motility features for which the simulations capture the full experimentally measured range; ideally, this number should be as close to the total number of motility features (26) as possible (**Figure 4G**). Finally, the mean feature overshoot indicates the average log10-ratio of the simulation range to the experimental range for each motility feature; ideally, we want this number to be greater than zero, but not much greater than zero, to prevent significant overshoot of the experimental motility feature distributions (**Figure 4H**). We reasoned that the ABM spatiotemporal resolution that best fits experimental data would maximize mean feature coverage ad number of covered features, while minimizing feature overshoot. A spatiotemporal resolution of 2 µm per voxel and a timestep size of 7.5 seconds (equivalent to a simulation run time of 77 timesteps to recapitulate the 9.5-minute experimental trajectories) satisfied these criteria.

### Parameterization of agent-based model to recapitulate each experimentally observed motility behavior

After calibrating the model spatiotemporal resolution, we sought to find which combinations of (MP, TP) recapitulate each experimentally observed motility behavior. The experimental data show that GCB and Tfh motility are heterogeneous and exploit seven different motility behaviors. Notably, each observed motility behavior is distinct from each other (i.e. each behavior has distinct speed and turning distributions), but each behavior also has within-group heterogeneity (i.e. a combination of different trajectories comprises each behavior). Therefore, to accurately model these behaviors, we need to implement a parametrization approach that can 1) simulate the observed between-group variance across behaviors and 2) simultaneously simulate the within-group heterogeneity of each behavior.

Because our goal is to ultimately use these simulations to predict how GCB and Tfh motility drive GCB-Tfh interactions, we implemented this parametrization approach in multi-cell simulations, in which cells are seeded at the expected density of the germinal center. In these multi-cell simulations, each cell agent contains additional volume exclusion rules to prevent cell-cell overlap (**Supplemental Text S3** & **Supplementary Figure 6**).

We first used multi-cell simulations to simulate several trajectories per (MP, TP) combination, with MP and TP ranging from 0 to 1. To assess which (MP, TP) values gave rise to simulated trajectories most similar to each experimental motility behavior, we computed the aggregated minimum Euclidean distance across seven orthogonal motility features (see *equation 1* in *Methods*) between the simulated trajectories generated by each (MP, TP) combination and the experimental trajectories that compose each experimental motility behavior (**Figure 4I**). This aggregated minimum distance computes the square root of the sum of squared Euclidean distances of the closest experimental trajectory to each simulated trajectory generated by an (MP, TP) combination (see *equation 1* in *Methods*). We found that for each experimental motility behavior, a unique subspace of MP and TP values yielded low aggregated minimum distances. This result suggests that distinct simulation parameter spaces can recapitulate each of the seven experimentally observed motility behaviors.

We next assessed whether the closest (MP, TP) combination to each motility behavior (i.e. the MP, TP pair with lowest aggregated minimum Euclidean distance to each experimental motility behavior) generated motility feature distributions similar to those of the experiment. We quantified the Cohen’s D parameter comparing simulated motility feature distributions and experimental feature distributions for each motility behavior (**Supplementary Figure 7A**). The Cohen’s D metric measures the relative standard deviation difference between two distributions, and a value less than or equal to 0.5 indicates a less than medium effect size. We found that the closest (MP, TP) combination in the single-cell simulation was highly dissimilar to experimental motility features (for each motility behavior, only about 50% of motility features had a Cohen’s D < 0.5). The speed and tortuosity features of each motility behavior were highly dissimilar between experimental and simulated groups (**Supplementary Figure 7A**). This result suggests that sampling a single (MP, TP) combination cannot effectively capture the distribution of all 26 motility features that we seek to recapitulate within each of the seven motility behaviors. Therefore, we sought to optimize a mixed distribution of several (MP, TP) combinations that can accurately recapitulate the distributions of experimental motility features for each motility behavior.

To derive this optimal mixed distribution of (MP, TP) values, we employed an evolutionary algorithm^36–38^ that iteratively optimized a combination of weights for each (MP, TP) combination, such that the group of simulated trajectories match those in each experimental motility behavior (see *Methods* and **Figure 4J**). We then compared simulated motility feature distributions generated from this mixed parameter distribution with the experimental data (**Figure 4K-N**, **Supplementary Figure 7B**). Similarity was significantly improved, with on average 86% of motility features having a Cohen’s D < 0.5 per motility behavior. Moreover, unlike in the prior optimization, the distributions for speed and tortuosity features are highly similar between experimental and simulated trajectories in each motility behaviors. Finally, we ran independent simulations that resampled the derived parameter distributions. These simulations show that re-sampling these distributions robustly recapitulate the same motility feature distributions per motility behavior (**Figure 4K-N**, **Supplementary Figure 7C**). Notably, our model recapitulates the within-group and between-group variance across speed and persistence features for each of the seven motility behaviors.

We compared top-down mixed distribution parameterization approach, described above, with a bottom-up, *k*-nearest neighbor approach (**Supplemental Text S4** and **Supplementary Figure 8**). The *k*-nearest neighbor approach computes a nearest-neighbor simulated trajectory for each experimental trajectory in each motility behavior. This approach then defines the (MP, TP) parameterization for each motility behavior as the distribution of (MP, TP) values that generated these simulated neighbor trajectories. We found that this approach was not robust; the (MP, TP) combination that yielded a single trajectory similar to an experimental trajectory did not consistently reproduce this trajectory across repeated simulations. Instead, our chosen top-down approach compares a population of stochastically generated trajectories from a given (MP, TP) combination to a set of experimentally measured trajectories within each motility behavior (rather than comparing individual pairs of trajectories). Therefore, the chosen mixed distribution approach was more robust because this approach accounts for the intrinsic variability across trajectories generated by a single (MP, TP) combination.

### Simulation predictions correlate with experimentally measured multi-cell interactions

After deriving parameter distributions that can recapitulate experimentally observed single-cell motility behaviors, we sought to validate that modeling these behaviors can recapitulate experimentally observed GCB-Tfh interactions.

Each of the 15 experimental movies in our intravital microscopy dataset had varying proportions of labelled cells (**Figure 5A**). Within each experiment, we can 1) measure interactions between these labelled cells and 2) validate whether simulations seeded with experiment-specific labeled cell density recapitulate these labelled interactions. Experimentally labelled cells were designated as either “tracked” cells or “segmented” cells. Tracked cells refer to cells labelled and segmented for 20 continuous frames, while segmented cells refer to any labelled cell that was segmented for at least 5 continuous frames. From each image set, we measured the GCB-Tfh contact time as the number of frames (out of 20 total frames) during which each tracked cell overlapped with any other segmented GCB or Tfh (see **Table 2** for details on interaction quantifications from the experiment and simulation). Because each experiment had different labelled volumes of GCBs and Tfhs, the experimentally measured GCB-Tfh contact time varied across experiments (**Figure 5B-C**). To validate our simulation, we ran simulations that recapitulated labelled cell volumes in each experiment and compared predicted labelled interactions with corresponding experimental measurements.

**Figure 5.**
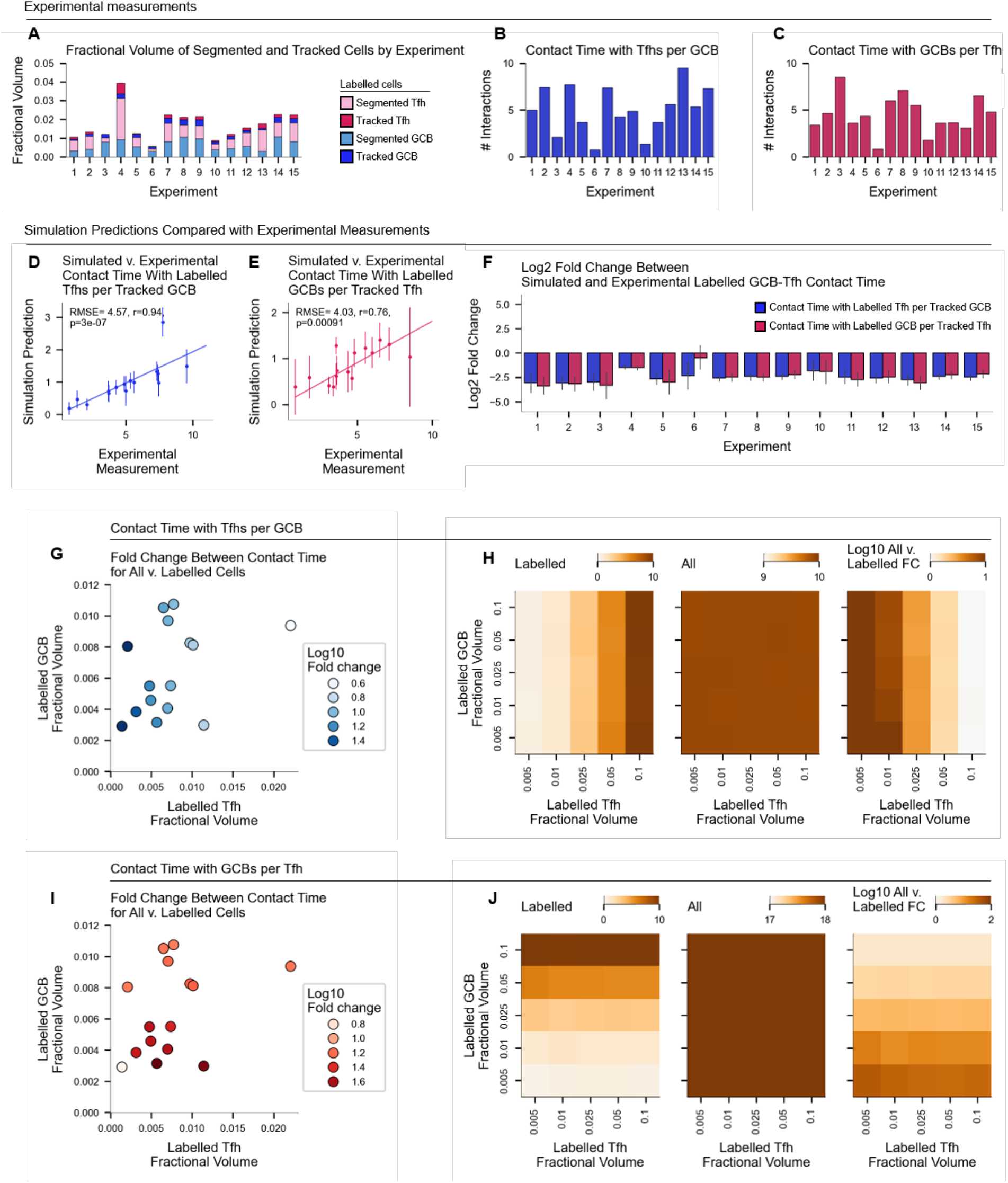
Validation of ABM predictions of GCB-Tfh interactions. **A,** Bar plot representing the fractional volume of segmented and tracked cell types within each intravital microscopy experiment. Segmented cells are cells that were tracked for 5 consecutive frames. Tracked cells are those that were tracked for 20 consecutive frames. **B-C,** Experimentally measured GCB-Tfh contact time in each intravital microscopy experiment. We measured contact time by calculating the volumetric overlap ratio between each tracked cell and any labelled cell type (which could have been tracked for 20 frames or segmented for a minimum of 5 frames; see Methods). **D,** Scatter plot of simulation predictions against experimental measurements for contact time with labelled Tfhs per tracked GCB. **E,** Scatter plots of simulation predictions against experimental measurements for contact time with labelled GCBs interactions per tracked Tfh. Vertical error bars represent the standard deviation from 100 simulation repetitions. The line indicates a linear regression with the intercept fixed at zero. ‘RMSE’: root mean squared error between average simulation predictions and experimental measurements, ‘r’: r-value for Spearman correlation, ‘p’: *p*-value for Spearman correlation. **F,** Bar plot representing the log_2_ fold change between simulation predictions and experimental measurements for contact time with labelled Tfhs per tracked GCB (blue) or labelled GCBs per tracked Tfh (red). Simulations were run at the experimental volume for 100 repetitions per experiment. **G,** Calculated fold change between predicted contact time with Tfhs per GCB for all versus labelled cells as a function of labelled Tfh and GCB volumes in each experiment. Each dot represents the log_10_-normalized average fold change from 100 simulation repetitions. **H,** A global parameter space exploration of how Tfh contact time per GCB varies for distinct combinations of labelled GCB and Tfh fractional volumes. Heatmaps show Tfh contact time per GCB across labelled cells, all cells, and the log_10_-normalized average fold change between all versus labelled contact time. Each simulation was run at the experimental volume for 50 repetitions. **I,** Calculated fold change between predicted contact time with GCBs per Tfh for all versus labelled cells as a function of labelled Tfh and GCB volumes in each experiment. Each dot represents the log_10_-normalized average fold change from 100 simulation repetitions. **J,** Parameter space exploration of how GCB contact time per Tfh varies for distinct combinations of labelled GCB and Tfh fractional volumes. Heatmaps show GCB contact time per Tfh across labelled cells, all cells, and the log_10_-normalized average fold change between all versus labelled contact time. Each simulation was run at the experimental volume for 50 repetitions. Multi-cell simulations represented in this figure were run with periodic boundary conditions at the experimental volume, a spatial resolution of 2µm per voxel, and temporal resolution of 7.5 seconds per timestep.

**Table 2.**
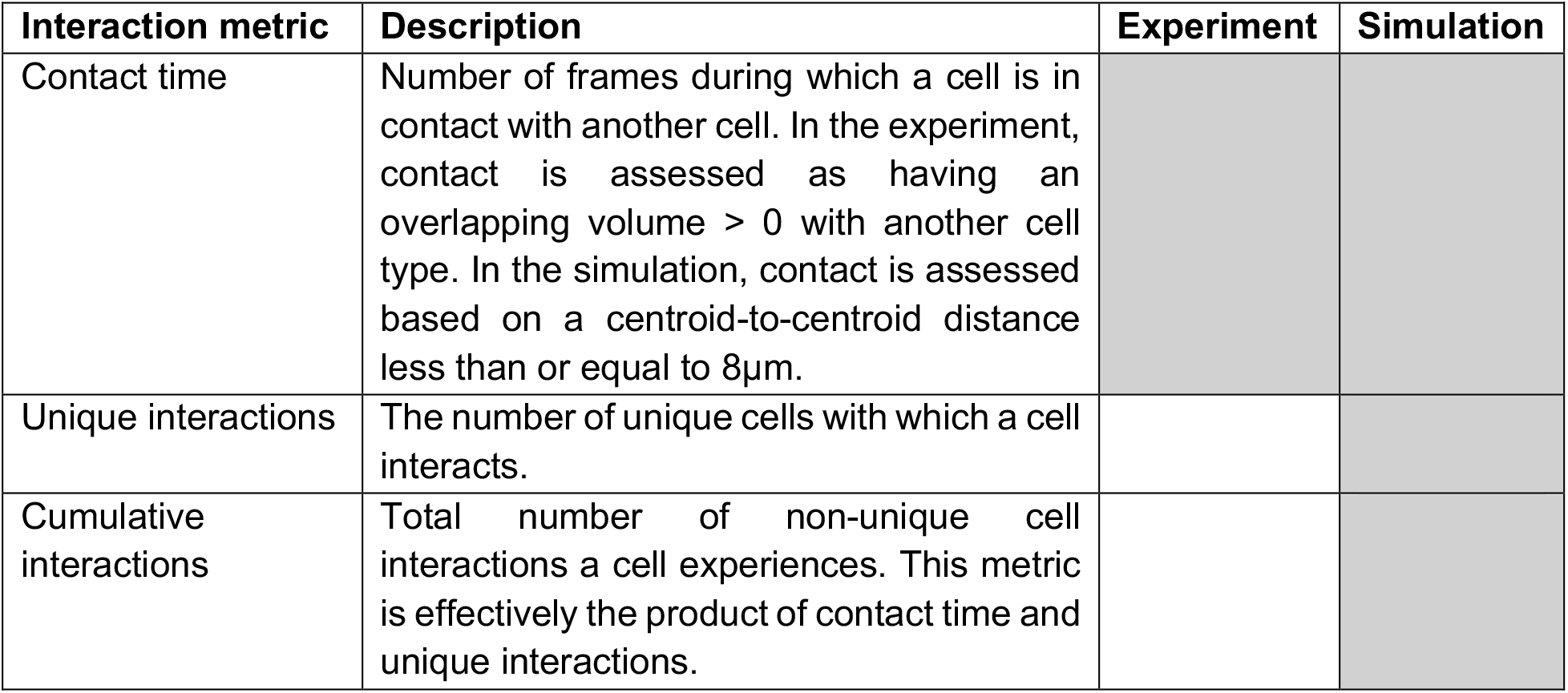
Interaction features computed within the experimental and simulation. In the experiment, contact time can only be computed between tracked and labelled cells. In the simulation all interaction metrics can be computed among tracked cells, labelled cells, or all cells.

| Interaction metric | Description | Experiment | Simulation |
|---|---|---|---|
| Contact time | Number of frames during which a cell is in contact with another cell. In the experiment, contact is assessed as having an overlapping volume > 0 with another cell type. In the simulation, contact is assessed based on a centroid-to-centroid distance less than or equal to 8μm. |  |  |
| Unique interactions | The number of unique cells with which a cell interacts. |  |  |
| Cumulative interactions | Total number of non-unique cell interactions a cell experiences. This metric is effectively the product of contact time and unique interactions. |  |  |

Within these validation simulations, we simulated the total expected GCBs and Tfhs in the GC volume and randomly selected a subset of these cells to be labelled according to experimental measurements (See **Supplemental Text S5** and **Supplementary Figure 9** for the calculation of labelled volumes and simulation runtime conditions). Based on prior literature, we assumed 40% of the simulated volume to comprise GCBs and 10% of the simulated volume to comprise Tfhs.^39^ Within these simulations, we modeled the populations of GCBs and Tfhs as moving according to weighted combinations of the seven motility behaviors that were experimentally observed (**Figure 3G**). According to this approach, the simulated GCB and Tfh populations were split across the observed occurrences of the seven behaviors (**Figure 3G**). For each simulated behavior, cell agents draw (MP, TP) values from the previously optimized mixed distributions for each motility behavior (**Figure 4J**). This approach allows us to model GCBs and Tfhs moving according to their experimentally observed, heterogeneous motility behaviors and then validate whether the emergent GCB-Tfh interactions that arise from these behaviors match experimental measurements.

We found that predictions for labelled Tfh contact time per tracked GCB were highly correlated with experimental measurements (R=0.94, *p* < 0.001, **Figure 5D**). Similarly, labelled GCB contact time per tracked Tfh was highly correlated with experimental measurements (R = 0.76, *p* <0.001, **Figure 5E**). Despite this high correlation, both simulations tend to underpredict experimental measurements by approximately a factor of four (**Figure 5F**). We hypothesize that our model underpredicts experimentally measured GCB-Tfh contact time because our model does not include additional rules for antibody-antigen affinity between GCBs and Tfhs. In the actual germinal center, GCBs and Tfhs maintain persistent contact, determined by antibody-antigen binding affinity, which could increase the experimentally measured GCB-Tfh contact time^6,7^.

After validating simulation predictions, we sought to apply simulations to understand how sparse labelling in the experimental imaging system impacts measured GCB-Tfh contact time. Recall that within the experimental data not all cells were labelled, and thus we could not measure GCB-Tfh contact time across all cells. However, our simulations were seeded with the total expected numbers of GCBs and Tfhs and we can, thus, compare predicted GCB-Tfh contact time across all cells with the contact time for the labelled subset of cells. We found that the fold change in contact time with Tfhs per GCB between all versus labelled cells ranged from ∼4 to 40-fold. Generally, as the labelled volume of Tfhs increased, the fold change between all and labelled contact time with Tfhs per GCB decreased (**Figure 5G**). To further characterize the difference between all and labelled contact time, we ran simulations for different labelled Tfh and GCB fractional volumes. In each simulation, we assumed 10% of labelled cells were tracked. Similarly, the fold change between all versus labelled contact time with Tfhs per GCB decreased as the labelled volume of Tfhs increased (**Figure 5H**). The fold change between contact time with GCBs per Tfh across all versus labelled cells ranged from ∼4 to ∼60-fold (**Figure 5I**). In simulations exploring different combinations of labelled Tfh and GCB volumes, we found that the predicted fold change between all and labelled contact time with GCBs per Tfh decreased, as the labelled GCB volume increased (**Figure 5J**). These results show that as the labelled cell volume increases and approaches the expected volume of each cell time, the difference between labelled measurements and all measurements of GCB-Tfh contact time decreased.

Finally, we were curious whether fast-moving cells were less likely to be measured in the experimental data because they have a higher likelihood of leaving the imaging volume (thus less likely to be tracked for 20 continuous frames). Our simulations show that while fast-moving cells are more likely to leave the imaging volume, adjusting for this measurement bias does not significantly impact predicted GCB-Tfh contact times of labelled cells (**Supplemental Text S6** and **Supplementary Figure 10**).

### Specific GCB and Tfh motility behaviors maximize unique GCB-Tfh interactions

After validating that simulations can recapitulate variations in experimentally measured GCB-Tfh contact time, we ran multi-cell simulations to systematically quantify how each experimentally observed single-cell motility behavior impacts unique GCB-Tfh interactions. In the previous section, we modeled GCB and Tfh populations as the heterogenous combination of behaviors 1-7 that were experimentally observed. Now, we sought to pinpoint how each of the individual behaviors influence GCB-Tfh interactions.

In our simulations, we can quantify unique interactions as the number of distinct cell agents a cell encounters (**Table 2**). We performed multi-cell simulations with different selected combinations of motility behaviors (B1-B7) for GCBs and Tfhs to predict the impact of motility behavior on unique GCB-Tfh interactions. (**Figure 6**). We ran these simulations in a (110 x 110 x 110) µm^3^ volume with static boundaries for 10 minutes, the duration of experimentally measured trajectories. The static boundaries mimic the experimental acquisition settings for imaging a static observation volume, in which cells are not tracked after leaving the imaging volume. To mimic these conditions in the simulation, when a cell agent leaves the simulation world, a new cell agent enters from the opposite face, instead of the same cell agent wrapping around the simulation world. We quantified cumulative interactions and unique interactions in these simulations (**Table 2**). Unique interactions indicate the total number of unique cell interaction partners a cell comes across. Cumulative interactions indicate the total number of cell-contacts a cell experiences (i.e. the product of unique cell interactions and length of persistent interaction with each interaction partner).

**Figure 6.**
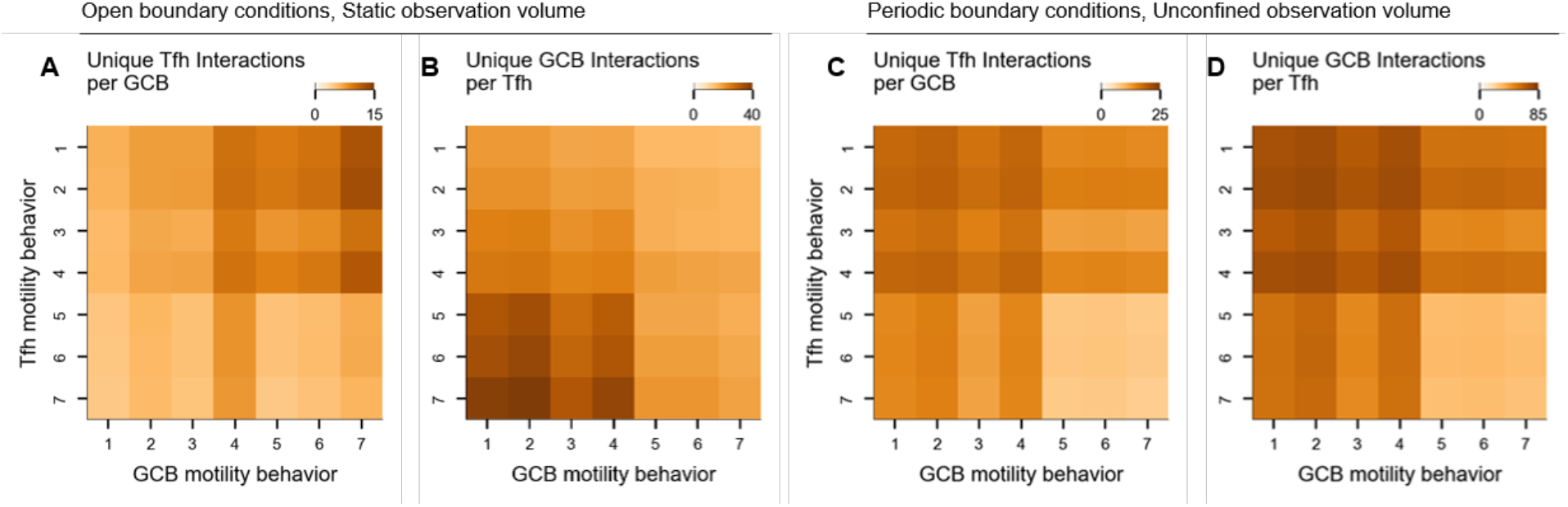
Application of ABM to predict how motility behavior drives unique GCB-Tfh interactions. **A-B,** Heatmaps representing **(A)** unique Tfh interactions per GCB and **(B)** unique GCB interactions per Tfh when running simulations of a static observation volume, in which Tfhs and GCBs follow each experimentally measured motility behavior (B1-B7). We simulated a static, constant-density observation volume by allowing cell agents to leave the simulation world and spawning new cells at the simulation boundaries whenever a cell agent left the world (see Methods). **C-D,** Heatmaps representing unique Tfh interactions per GCB **(C)** and unique GCB interactions per Tfh **(D)** when running simulations with periodic boundary conditions in which Tfhs and GCBs follow each experimentally measured motility behavior (B1-B7). These simulations allowed cell agents to leave the simulation world and re-enter as the same cell agent (see Methods). Each simulation volume was 40% occupied by GCBs and 10% occupied by Tfhs. The simulation world dimensions were 110 x 110 x 110 µm (e.g., 55 x 55 x 55 voxels). All simulations were repeated 50 times.

We found that out of all motility behavior combinations, Tfhs moving according to motility behaviors B1-B4 and GCBs moving according to motility behaviors B5-B7 yielded the maximum unique Tfh interactions per GCB (**Figure 6A**). This result is interesting because this breakdown reflects the baseline motility behaviors of GCBs and Tfhs measured experimentally (**Figure 3G**). Conversely, Tfhs moving according to B5-B7 and GCBs moving according to B1-4 maximizes unique GCB interactions per Tfh (**Figure 6B**). A major driving factor of this observed result is that slow-moving cells (e.g. B5-B7) are less likely to leave the simulated volume and spend more time on average within the simulation volume than fast-moving cells (**Supplementary Figure 11A**). Thus, when we normalize the unique Tfh interactions against the number of simulated GCBs, we predict a higher normalized count per cell because there are fewer GCBs to normalize against (**Supplementary Figure 11B**). These results suggest that the predicted high unique Tfh interactions observed per GCB in a static observation volume when GCBs move slowly are a result of slow-moving GCBs having a higher ability to stay in the observation volume than faster-moving counterparts.

We also ran this parameter-space exploration in a periodic volume, in which cells wrap around the simulation and retain their identities. While static boundary conditions represent the conditions of experimental observations well, periodic boundary conditions better represent what a given fast-moving individual cell ‘sees’ in an unconfined volume. With periodic boundary conditions, the trend is slightly different: both GCBs and Tfhs moving according to fast motility behaviors (B1-B4) yielded the highest number of unique GCB-Tfh interactions (**Figure 6C-D**). This trend arises because, in an unconfined volume, moving as fast as possible allows both cells to surveil the maximum possible space and engage in cell-cell interactions. Altogether, these results suggest that GCBs can experience comparable interactions with Tfhs without having to move as fast, allowing GCBs to conserve energy for other processes.

Taken together, these simulations suggest that, within a static observation volume, slow-moving GCBs can remain in place and maximize their ability to encounter fast-moving Tfhs, which matches the baseline observed motility behaviors of these cells.

## DISCUSSION

In this study, we present PRISMM, a data-driven and mechanistic approach to quantify heterogeneous lymphocyte motility behaviors and predict how these behaviors contribute to the likelihood of cell-cell interaction. Our pipeline uses unsupervised clustering to identify single-cell motility behaviors from *in vivo* lymphocyte trajectories. PRISMM then uses 3D agent-based simulations to mechanistically predict how these motility behaviors influence the likelihood of GCB-Tfh interactions. Using PRISMM, we identified seven distinct lymphocyte motility behaviors with characteristic speeds and tortuosities. We found that Tfhs tend to occupy motility behaviors characterized by high cell speeds, while GCBs were abundant in low-speed motility behaviors. Our agent-based model predicts that this division of motility behaviors maximizes unique GCB-Tfh interactions within static observation volumes.

While prior literatures has shown that shifts in baseline GCB and Tfh motility behaviors can impair the antibody response,^4,5^ prior work has not directly mapped perturbations of lymphocyte motility behaviors to the quantity of GCB-Tfh interactions. The relationship between lymphocyte motility and cell-cell interaction has been difficult to quantify using intravital imgaing, because sparse labeling for single-cell segmentation obscures GCB-Tfh interactions across unlabeled cells within the system. Using our simulation, we modeled the expected total numbers of GCBs and Tfhs within the GC and predicted the full extent of cumulative and unique GCB-Tfh interactions amongst this complete population. Using our model, we can elucidate the difference between predictions of GCB-Tfh interactions across all cells versus the labelled subset (e.g. we quantified that the predicted number of Tfh contacts that a GCB experiences was on average 12 times higher than that observed between labeled cells in our experiments).

In addition to measuring the difference between unlabelled and labelled interactions, our model can systematically quantify how distinct motility behaviors influence unique GCB-Tfh interactions. We found that within a static observation volume, the number of unique Tfh interactions per GCB was predicted to be highest when Tfhs moved according to motility behaviors B1-B4 and GCBs moved according to motility behaviors B5-B7. Using periodic boundary conditions, we found that slow-moving GCBs and fast-moving GCBs experience similar numbers of unique interactions with fast-moving Tfhs. These predictions suggest that the baseline motility behaviors of GCBs and Tfhs enable these cell types to efficiently surveil the germinal center environment (**Figure 6**). Our model further suggests that dysregulation of Tfh motility towards slower motility behaviors can reduce the number of unique Tfh interactions experienced per GCB, providing a mechanism by which dysregulated cell motility may lead to diminished immune responses that has been observed in other experimental studies^4,5^.

A known limitation of intravital imaging studies is that measured motility behaviors are biased towards trajectories that start and end within the imaging volume. This constraint could bias measurements of slow-moving trajectories. Using our simulation, we predicted the likelihood of a given motility behavior escaping the imaging volume and used these likelihoods to adjust the proportion of observed motility behaviors (**Supplementary Figure 10**). We then considered whether this adjustment significantly impacts predicted GCB-Tfh interactions. Our results show that despite experimental bias towards undercounting the abundance of fast-moving motility behaviors, adjusting for this undercounting does not significantly impact predicted interactions (**Supplementary Figure 10F-G**). This result suggests that while intravital imaging can undercount fast-moving behaviors, the emergent impact of this bias was not substantial in this study.

Prior studies have leveraged computational modeling to understand how immune cell motility enables cell-cell interactions. Many prior models simulate immune cell motility as a persistent random walk^40–43^ in which the average speed and persistence time (average time before the cell significantly changes direction) of lymphocytes were approximated in bulk from tracked trajectories seen in *in vivo* intravital microscopy videos. Here, we go further by quantifying several distinct motility behaviors from *in vivo* intravital microscopy data and then identifying mechanistic parameters that can recapitulate each of these behaviors. This approach allows us to model the known heterogeneity of cell motility behaviors^44^ by modeling a single cell type as a collection of distinct, experimentally observed motility behaviors. Moreover, our stochastic model simulates the intrinsic variability within a single motility behavior. By using a distribution of probabilistic movement and turn probabilities, we can simulate a population of stochastic trajectories that mimic the heterogeneity in the population of experimentally measured trajectories. In this way, this pipeline provides a natural extension of prior modeling efforts, allowing us to probe how heterogeneous motility patterns of B-cells and T-cells influence their ability to engage in cell-cell interactions that carry out the adaptive immune response.

While our simulation provides key insights about how GCB and Tfh motility impact GCB-Tfh interactions, our model currently does not capture other important cellular mechanisms within GC biology. Our simulation predictions of GCB-Tfh contact time are highly correlated with experimental measurements but still underpredict this metric by a factor of 4. We hypothesize that this underprediction stems from the fact that our current model does not simulate important GC mechanisms such as affinity-driven GCB-Tfh interactions and GCB-Tfh co-localization through chemotaxis. These mechanisms could prolong predicted GCB-Tfh contact time and better match experimental measurements. Adding these rules to the simulation would not be a trivial addition; increasing interaction time would further affect trajectories and, in turn, simulated motility patterns, requiring further refinement of the optimization of model movement parameters. Several prior studies have modelled the interaction dynamics and spatial orientation of the germinal center reaction^31,45–49^. Future work could integrate our model of heterogeneous cell motility with these computational models to elucidate further how distinct lymphocyte motility behaviors drive varied immune responses. Extensions of this work could also consider how aging or genetic mutations that dysregulate lymphocyte motility emergently impact GCB-Tfh interactions during the germinal center response.^50–54^

Cell motility drives cell-cell interactions that are critical for tissue function in several biological systems. PRISMM lays the foundation to understand how heterogeneous motility behaviors drive interactions across a variety of tissue systems under both physiological and pathophysiological conditions.

## DATA & CODE AVAILABILITY

Cell motility data will be deposited to our Github repository, which will be publicly available upon manuscript publication. Original code developed for this manuscript is available on our Github repository, which will be publicly available upon manuscript publication. The link to our Github repository can be found here: https://github.com/nikita-sivakumar/prismm

## ACKNOWLEDGMENTS

We acknowledge the financial support for this work from the National Science Foundation Graduate Research Fellowship Program, Metro Washington Chapter Achievement Rewards for College Scientists (ARCS) Fellowship, NTT Research Fellowship, and Siebel fellowship (to N.S.); the National Institutes of Health under award numbers R35GM157099 (to J.M.P.); and the 2024 Salisbury Family and Center for Innovative Medicine Human Aging Project (HAP) Scholar Award (to J.M.P.). W.B. was supported by NIH NCI R01 CA270245, LLS TRP 6641-22, LLS TRP 6679-24, FLF CURE FL 224362, LRF FL PRG 226504, ASH Junior Faculty Scholar Award 202414 and 223390, and Gilead Sciences Research Scholars Program GSI 231647-01.

## AUTHOR CONTRIBUTIONS

N.S., J.M.P., and F.M.G. conceived this study. K.C., C.X., and W.B. performed imaging and animal experiments, collected data, and contributed reagents. N.S., C.M., J.M.P., and F.M.G. conceived data analysis. N.S. and C.M. performed formal analysis. All authors interpreted the results. N.S., J.M.P., and F.M.G. wrote the manuscript. All authors contributed to reviewing and editing the manuscript. J.M.P and F.M.G. supervised the study. W.B., J.M.P. and F.M.G. contributed equally as co-corresponding authors.

## SUPPLEMENTAL INFORMATION

Document S1. Supplemental Text S1-S6, Supplementary Table 1, Supplementary Figures 1-12.

## METHOD DETAILS

We developed a data-driven simulation pipeline to 1) identify emergent cell motility behaviors from three-dimensional (3D), *in vivo* intravital time-lapse microscopy data and 2) simulate these single-cell motility behaviors within multi-cell simulations to predict how differential motility behaviors drive the likelihood of cell-cell interactions. Below, we discuss this pipeline in two parts. First, we discuss the experimental imaging setup and the data-driven analysis of experimental data to identify emergent motility behaviors that characterize single-cell trajectories. Then, we discuss the process of building and validating agent-based models (ABMs) to recapitulate single-cell motility behaviors within 3D volumes and predict emergent cell-cell interactions.

### Mouse intravital microscopy system to image GCB and Tfh cell motility

To visualize lymphocytes within mouse germinal centers, we transplanted bone marrow cells from R26-CFP;Cγ1-cre or R26R-EYFP;Cγ1-cre and R26R-tdTomato;CD4-cre donor mice into recipient mice. After immune cell repopulation in recipient mice, germinal center (GC) responses were initiated using standard procedures, with NP(4-hydroxy-3-nitrophenylacetate)-ovalbumin (2 mg/ml in PBS, N-5051, Biosearch Technologies) mixed with alum at 1:1 volume ratio (total 50 µl) injected into the footpad. Two-photon intravital imaging of the cells within the popliteal lymph node was performed at the peak of the GC reaction between days 9 and 11 (see **Supplementary Table 1** for further imaging conditions). The image acquisition interval was 30-seconds and the imaging duration ranged between 80 to 90 minutes. The imaging volume was 231 µm x 231 µm x 90 µm. A subset of the tracked GCBs in the imaged mice carried an EZH2 (Y641F) mutation. Because the main purpose of this study was to quantify differences between Tfh and GCB motility, we combined wild-type and mutant GCBs into a single GCB population for analysis. While wild-type and mutant-EZH2 GCBs have distinct motility behaviors,^51^ their observed behaviors were both distinct from that of Tfhs (**Supplementary Figure 12**). Combining wild-type and mutant trajectories enables us to sample the universe of possible GCB motility behaviors, while increasing the number of observations for the GCB population. Throughout the text, when referring to the “GCBs” we are describing this combined wild-type and mutant-EZH2 population. Moreover, in our simulations, we model the combined GCB population, capturing the motility and emergent impact of both wild-type and mutant motility behaviors, which is appropriate as we are comparing the results of these simulations to experiments in which both wild-type and mutant cells are present. More detail on the differences between WT-EZH2 and MT-EZH2 GCBs were reported in other studies,^51–54^ and these differences were not the focus of the present study.

#### Bone marrow transplant

Cγ1-cre mice (#010611, The Jackson Laboratory) were crossed with R26-CFP^55^ or R26R-EYFP mice (#006148, The Jackson Laboratory) to obtain GCBs fluorescently labelled with either CFP or YFP. CD4-cre mice (#022071, The Jackson Laboratory) were mated with R26R-tdTomato mice (#007909, The Jackson Laboratory) to generate Tfhs fluorescently labelled with tdTomato. Bone marrow cells were harvested from these donors at 8-12 weeks and mixed with bone marrow cells from Cγ1-cre colorless donors. Harvested cells were injected intravenously into C57BL/6J host female mice (#000664, The Jackson Laboratory). Recipient mice were lethally irradiated with 450 rads the day before and an additional 450 rads 2 hours prior to transplantation.

All animal experimentation and housing procedures complied with institutional guidelines established by Cornell University Institutional Animal Care and Use Committee, the Weill Cornell Medical College, the Guide for the Care and Use of Laboratory Animals (National Academy of Sciences 1996), and the Association for Assessment and Accreditation of Laboratory Animal Care International. Mice were maintained in the Cornell University animal facility under conventional housing conditions (22.3–22.7°C; 38–40% humidity; 12-h light/dark cycle) with ad libitum access to food and water.

#### Intravital two-photon microscopy

We performed intravital imaging of mouse popliteal lymph node GCs four to six months after bone marrow transplant. We prepared the popliteal lymph node for imaging based on previously established methods^18^. Imaged mice received 1-1.5% isoflurane in oxygen for anesthetization. Before imaging, we shaved the leg skin using a shaver and hair removal cream (Nair). Using microdissection forceps, we made an incision behind the knee and removed fatty tissue covering the popliteal lymph node. We then hydrated the popliteal lymph node with saline and embedded the lymph node within in a cover glass. We applied grease below the cover glass to prevent saline evaporation. We placed a heating wire (nichrome) above the cover glass to maintain lymph node temperature at 36.5 ± 0.5 °C. We used a temperature probe (IT-23, Braintree Scientific) below the cover glass to monitor lymph node temperature. We maintained the mouse core body temperature at 36–37 °C using a temperature controller (FHC) comprising a rectal probe and a heating pad.

We used a two-photon microscope (Bergamo II with ThorImage 4.1, Thorlabs) with a NA 1.05 objective (XLPLN25XWMP2, Olympus) for image acquisition. We used a mode-locked Ti:sapphire laser (Chameleon Vision-S, Coherent) as the excitation source. We used several band-pass filters to image CFP, YFP, and tdTomato simultaneously (FF01-475/42, FF03-525/50, FF01-585/29, Semrock). The imaging volume was 231 µm x 231 µm x 90 µm. We acquired images at 30-second intervals.

### Data-driven analysis of trajectories from intravital time-lapse microscopy images

The overall pipeline for data-driven analysis of the intravital microscopy dataset includes 4 steps (**Figure 2**). First, we used the commercial software IMARIS (Oxford Instruments) to segment and track single-cell trajectories in each time-lapse image set. Second, we computed 36 motility features from each trajectory to quantify displacements and angular characteristics for each cell (**Table 1** and **Supplemental Text S1**). Third, we applied factor analysis to identify orthogonal features that differentiated patterns of trajectories within the dataset, and unsupervised clustering to define distinct motility behaviors. Finally, to improve quality and reduce noise, we eliminated artifactual trajectories based on outlier feature values. We repeated clustering on the cleaned dataset (**Supplemental Text S1**). We expand upon each of these steps in detail in the following sub-sections.

#### Extraction of trajectories from intravital time-lapse microscopy images

To extract single-cell trajectories from each intravital time-lapse microscopy image set, IMARIS software was used to segment cells and track individual trajectories. A total of 9,720 cell trajectories were extracted from the videos of the 15 mice. We tracked trajectories for 9.5 minutes (20 frames) and, for consistency in comparison, analyzed only nonoverlapping complete 20-frame sequences. We also segmented cells within the imaging volume for at least 2 minutes (5 frames) to estimate the density of segmented cells. Throughout the text, we refer to “tracked” cells as those continuously tracked for 20 frames and for which we derived a cell trajectory. We refer to “segmented” cells as those that were tracked continuously for at least 5 frames and were a part of the “labelled” cell population measured within the image, but not necessarily used to derive a motility trajectory. In IMARIS, we tracked the overlapping surface-to-surface volume of tracked cells with any segmented cells over time. We processed this metric to derive a binary state of cell-cell interaction, such that if the overlapping volume (continuous value) between two cells was greater than zero those two cells were considered to be interacting for that frame (binary state). From this metric, we derived the total contact time with Tfhs and GCBs of each cell (see *Comparison of simulation interactions with experimental measurements* below).

#### Motility feature extraction from each trajectory

For each of the 9,720 trajectories, we extracted 36 motility features to describe each trajectory (Table 1). These features describe the displacement and angular displacement aspects of the trajectory. To further characterize directional biases in each trajectory, we applied singular value decomposition to project the 3D trajectory into primary, secondary, and tertiary axes that explain decreasing levels of variance. Displacement distribution features were measured in the primary, secondary, and tertiary axes. To reduce the feature space prior to dimensionality reduction and unsupervised clustering, we applied factor analysis. We performed factor analysis to derive a communality score that describes how much variance each individual features explains within the data. We selected features with a communality score > 0.5 for behavior phenotyping.

#### Dimensionality reduction & unsupervised clustering of single-cell trajectories

For ease of visualization, we projected the motility features for each trajectory onto a two-dimensional UMAP embedding. To characterize how this UMAP embedding differentiated trajectories, we colored the UMAP space according to the value of each motility feature. We then applied unsupervised hierarchical clustering and *k*-means clustering to motility features (selected by factor analysis, as described in the previous section) to identify similar groups of cells/trajectories within the data. To phenotype each cluster, we plotted the distribution of key motility features across each cluster. In this process, we also identified clusters with outlier values for certain motility features and re-examined trajectories within these clusters to assess whether they were appropriately segmented/tracked. Based on this process, we developed rules to eliminate outlier trajectories. We repeated the steps of motility feature extraction, dimensionality reduction, and unsupervised clustering to identify and classify cells by discrete motility behaviors (see *Results* section and **Supplemental Text S1**). The cleaned dataset contained 6.912 trajectories.

### Simulation overview

We developed a three-dimensional (3D) on-lattice agent-based model (ABM) to simulate single-cell motility behaviors and multicellular interactions. Cells are simulated as autonomous agents that evaluate a probabilistic decision to move at each simulated timestep in the ABM. Our on-lattice assumption assumes that the centers of cells can only move between the centers of neighboring voxels in our grid-based simulated environment. We conducted both single-cell and multi-cell simulations. Within single-cell simulations, cell agents move independently of environmental obstructions. In multi-cell simulations, cell agents follow additional rules governing contact and overlap with surrounding cells. Below, we discuss the cell-agent movement rule that governs individual cell movement decisions on each simulated timestep within the ABM environment. Then, we discuss nuances in the simulation environment and ruleset for single-cell and multi-cell simulations.

#### Cell agent movement rule

At each timestep, cell agents execute a motility rule governed by two parameters: the movement probability (MP) defines the probability that the cell moves to a neighboring voxel in the simulation space during one timestep; and the turn probability (TP) dictates whether the cell changes its heading during this movement. If the turn probability is not satisfied, the cell will move to the voxel it is already pointing to — i.e., it will move persistently. If the turn probability is satisfied, the cell will change direction before moving. Because turn behaviors only emerge when the cell decides to move, the effective likelihood of a cell turning is the product of MP and TP.

#### Spatiotemporal resolution of simulations

The experimental data we sought to recapitulate consisted of 3D single-cell trajectories lasting 9.5 minutes and collected over 20 consecutive frames (i.e., a temporal resolution of 30 seconds and 19 possible cell movements). In our models, we had to determine the spatiotemporal resolution that best recapitulated the motility properties of the experimental data. The simplest solution for cellular ABMs is typically to assume a voxel size equal to the cell volume. But here, that assumption would confound the resolution of simulated trajectories, because a cell could only move a cell-size distance per timestep. In contrast, in the experimental data, cells may move any distance within a continuous range between frames. If simulated cells can only move one cell-size distance at a time, the simulated trajectories may not capture the fine-grained motility behaviors of cells in our system.

To enable our simulations to capture the behavior of both slow- and fast-moving cells, we tested voxel sizes of 1-4 µm and timestep sizes of 5-30 seconds, while keeping the overall simulation duration at 9.5 minutes. For each spatiotemporal resolution pair (voxel size, timestep size), we generated a simulated dataset of trajectories by co-varying MP and TP from 0 to 1. We then benchmarked the simulated trajectories against the experimental data by comparing the distributions of several motility features extracted from each trajectory. Based on this analysis, we chose a spatial resolution of 2 µm and a temporal resolution of 7.5 seconds (i.e., 76 time-intervals or 77 timesteps), because this spatiotemporal resolution captured the full range of experimental motility parameter distributions.

Our simulation used a voxel size of 8µm^3^ and a timestep of 7.5 seconds, and we assumed that all cells had a uniform radius of 4µm. The simulation voxel size was smaller than the cell size. In multi-cell simulations we applied volume-exclusion rules to allow cells to interact without occupying the same space (see ***multi-cell simulations*** below). To simulate the intercalation of cells as they move within a 3D environment, we simulated cells with an effective minimum radius of 2µm, allowing them to overlap up to 2µm. This overlap simulates the deformability of cells as they contact and interact with each other.

#### Single-cell simulations

In single-cell simulations, cell agents evaluate MP and TP to move through the simulation environment without considering environmental obstructions; i.e., cells do not follow additional rules to prevent overlap with surrounding cells. Because cells do not need to apply volume-exclusion rules (unlike *multi-cell simulations;* see below), single-cell simulations are more efficient to run. Therefore, we initially used single-cell simulations to assess how MP and TP affect the motility features of simulated trajectories. We then compared simulated trajectories from single-cell simulations with those from multi-cell simulations to evaluate whether single-cell simulations can be used to fit model parameters (i.e., MP and TP) to recapitulate experimentally measured motility behaviors.

In single-cell simulations, cells are seeded at the center of the simulation world with a random heading. The simulated world size is set sufficiently large so that the cell cannot escape the simulated environment during runtime. For example, if the simulation run time was 20 timesteps, the simulated world size spanned −20 to 20 voxels in the x, y, and z directions. Each simulation was run with a constant MP *and* TP value, using 500 cells to generate 500 trajectories per (MP, TP) combination.

#### Multi-cell simulations

The multi-cell simulations model multiple cellular agents within a simulation world. These simulations enable us to predict how cell density can affect cell trajectories. Moreover, this framework allows the model to detect cells that are in contact (interacting). Unless otherwise noted, multi-cell simulations comprised 50% of the simulated volume occupied by GCBs and Tfhs, as observed in prior literature.^39^

The multi-cell simulations include additional criteria to the cell movement rule to prevent cell-cell overlap (**Supplementary Figure 6A**). If a cell decides to move persistently and the voxel ahead is occupied, the cell searches for an empty voxel that neighbors both its current location and the neighboring voxel it is pointing to. If any of these voxels are unoccupied, the cell moves to one of them at random with equal likelihood; if all voxels are occupied, no movement occurs. This movement behavior allows cells to navigate around obstacles while maintaining a degree of persistence. This consideration is unnecessary for a cell when it is turning, since the cell already has several voxels to choose from that might be unoccupied. In multi-cell simulations, cells are allowed up to 2µm of overlap with each other to simulate the ability of cells to intercalate as they move.

In multi-cell simulations, cells are seeded at random locations in the simulation world with a random directional heading. Multi-cell simulations assume that we are simulating a representative subspace of the entire experimental system; accordingly, they apply periodic boundary conditions. Cells are allowed to wrap around the simulation world and retain their identity. This wrapping behavior assumes that the simulated space is a sub-volume of a larger, relatively homogeneous environment and, therefore, that if a cell left the simulated space, it would encounter an environment effectively similar to the one we are simulating. This assumption allows us to track individual cellular interactions throughout the simulation. This assumption also allows us to save computational resources by simulating a smaller volume than the experimental imaging volume. We run select simulations at the exact imaging volume to check that the smaller-world simulations are consistent. In addition, we ran some multi-cell simulations with the assumption that the simulated space represented a static observation volume. These simulations employed static boundary conditions, meaning that when a cell left the imaging volume, a new cell was spawned at the wrapped location to maintain cell density within the simulation. We note in the results whether we had used static or periodic boundary conditions and how to interpret the simulations accordingly. Static observation volume simulations depict conditions in the experimental imaging systems.

### Optimization of simulation motility parameters to recapitulate experimental motility behaviors

We sought to parameterize our simulation to the distinct motility patterns identified in the experimental data using unsupervised clustering. We first generated a “simulated trajectory space” by running simulations over different values of MP and TP. We sampled MP uniformly from 0.05 to 0.95 (19 unique MP values) and we sampled TP from 0 to 1, including higher resolution at small values ([0, 0.0005, 0.001, 0.005, 0.01, 0.025, 0.05, 0.1, 0.2, 0.3, 0.4, 0.5, 0.6, 0.7, 0.8, 0.9]; 16 unique TP values).

We then assessed how similar each (MP, TP) parameter combination was to each experimentally measured motility behavior, by quantifying the aggregated minimum Euclidean distance between simulated trajectories in each (MP, TP) parameter combination and the experimental trajectories within each experimental motility behavior. This Euclidean distance was evaluated for seven orthogonal features in the motility feature space (average speed, MSD (τ=1 minute), MSD (τ=30 seconds) in the secondary axis, MSD (τ=30 seconds) in the tertiary axis, average angular displacement, progressivity, and net distance). This metric describes how similar the simulated trajectories generated by a given (MP, TP) combination are to experimentally observed trajectories in a given motility behavior. This distance metric is as follows:

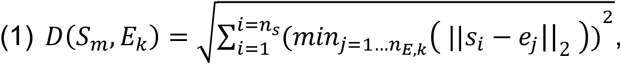

Where *S_m_* is the set of simulated trajectories in a single (MP, TP) combination, *E_k_* is the set of experimentally observed trajectories in motility behavior *k*, *n_s_* is the number of simulated trajectories and *n_E_*_,*k*_ is the number of experimental trajectories.

We then used an evolutionary algorithm to find the optimal mixture of (MP, TP) pairs that recapitulate each experimentally observed motility behavior. We fed in (MP, TP) pairs with minimum aggregated distances within the 75^th^ percentile of the overall minimum aggregated distance to the evolutionary algorithm to exclude highly dissimilar parameter sets. We chose to use an evolutionary algorithm, because this approach can efficiently search high-dimensionality parameter spaces of stochastic models.^37,38^ The evolutionary algorithm first evaluated a set of initial guesses for the mixed distribution and iteratively improved them by recombining choices with high similarity to the experimental data. The evolutionary algorithm specifically aims to minimize a loss function that computes the mean squared difference between the experimental and simulated cumulative probability distributions for a subset of four motility features (MSD (τ=1 minute), average angular displacement, progressivity, and net distance). The loss function computed for the minimal set of features *f* is as follows:

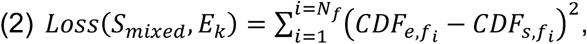

Where *S_mixed_*is the set of simulated trajectories sampled from the optimized mixed (MP, TP) distribution, *E_k_*is the set of experimentally observed trajectories in motility behavior *k*, and CDF represents the empirical cumulative density function for motility feature *f*. We tested several loss functions and found that minimizing the differences between CDFs led to the best convergence towards a solution.

Taken together, this approach allows us to derive parameter distributions that give rise to a simulated set of trajectories that match the heterogeneity of an observed group of trajectories that we want to recapitulate.

### Quantification of simulated interaction dynamics

We extracted several features in the multi-cell ABM to characterize interaction dynamics (**Table 2**). Assuming that simulated cells have a radius of 4 µm, we defined interacting cells as those with a centroid-to-centroid distance less than or equal to 8 µm. For each cell agent, we computed contact time with GCBs or Tfhs as the number of timesteps that a cell agent is within 8 µm of any GCB or any Tfh. This metric is directly comparable with experimental measurements (see *Comparison of simulation interactions with experimental measurements* below). In addition to cell-cell contact time, we computed each cell agent’s number of unique interactions and cumulative interactions. Unique interactions count the number of distinct cell interaction partners that a cell agent encounters. Cumulative interactions count all interactions across all timesteps (i.e., cumulative interactions could include interactions with the same cell that occur over multiple timesteps).

### Comparison of simulation interactions with experimental measurements

To validate simulation predictions of interactions, we derived metrics from the simulation that were directly comparable with experimental measurements. First, we ran simulations that recapitulated the labelled Tfh and GCB fractional volumes from each experiment. Then, we compared simulation predictions of time in contact with GCBs or Tfhs per cell type with corresponding experimental measurements from each experiment.

#### Simulating labelled volumes in each experiment

To compute the labelled fractional volume of GCBs and Tfhs in each experiment, we divided the product of the number of labelled cells and cell volume by the entire imaging volume. We assumed each cell had a radius of 4µm and, thus, a spherical volume of approximately 268µm^3^. We repeated this method to compute the fractional volume of segmented cells and tracked cells for GCBs and Tfhs in each experiment. Then, we ran simulations with the expected density of GCBs and Tfhs derived from literature (0.4 and 0.1 fractional volume, respectively). We then randomly selected simulated GCBs and Tfhs as “segmented” or “tracked” on their respective fractional volumes in each experiment.

#### Comparing simulation predictions with experimental measurements

In each experiment, we derived the proportion of each tracked cell that overlapped with a GCB or Tfh. For each frame in which this proportion was greater than zero, we considered the tracked cell to be in contact with a GCB or a Tfh. Similarly, in the simulation, we computed the number of timesteps at which each tracked GCB and tracked Tfh cell agent overlapped with a labelled GCB or labelled Tfh agent. Because we ran the simulation four times as long as the experiment, we divided this metric by four to make it comparable to the experimental measurements. Finally, we compare the contact time with Tfhs per GCB and contact time with GCBs per Tfh with corresponding experimental measurements from each experiment.

