## Supplemental materials for "Data-driven simulations elucidate how lymphocyte motility behaviors influence cell-cell interactions within germinal centers"

This file contains:

- Supplemental Text S1-S6
- Supplementary Table 1
- Supplementary Figures 1-12

**SUPPLEMENTAL TEXT S1. Initial clustering of lymphocyte trajectories indicated outliers.** To quantify behavioral phenotypes across the experimentally-obtained lymphocyte trajectories, we extracted 36 motility features from each trajectory. These features described the overall displacement and angular displacement properties of the trajectory (**Figure 2, Table 1**). We then performed feature selection and unsupervised clustering to identify "clusters" of trajectories with emergently similar motility feature values. To select features for unsupervised clustering, we ran a factor analysis to reduce the total features considered in the analysis to orthogonal features that explain high variance in the data. PCA showed that 5 PCs explain ~75% of the variance (**Supplementary Figure 1A**). Factor analysis on these 5 PCs gave communality scores that describe the degree of variance each feature explains across the 5 PCs. We selected features with communality scores > 0.5 for further analysis. This process eliminated 5 features. The eliminated features included shape properties of the angular distribution and progressivity in non-primary axes of movement. Ultimately, the reduced feature space contained three main groups of features (**Supplementary figure 1B**): net distance/progressivity in 3D and primary axis of movement; shape features of the displacement distribution in the primary and secondary axis; and average speed and MSD features in all axes of movement.

For ease of visualization, we constructed a two-dimensional (2D) uniform manifold approximation and projection (UMAP) embedding of the 9,720 trajectories as described by 31 motility features. To extract patterns of motility features from this high-dimensional data, we then performed unsupervised *k*-means clustering (**Supplementary Figure 1C**). Within this initial clustering, we found that speed and tortuosity were key differentiators of motility behaviors (**Supplementary Figure 1D**). We found that Tfhs generally exhibited high-speed motility behaviors, while GCBs tended towards low-speed motility behaviors (**Supplementary Figure 1E**).

Within the initial clustering of the entire dataset, we found that one cluster predominantly contained trajectories with high displacement kurtosis and skewness values (**Supplementary Figure 1F**). Examination of the displacement distribution of individual trajectories within this cluster indicated unusually high single-frame displacements, while the displacement distribution of trajectories within other clusters of the data did not contain this property (**Supplementary Figure 2A**). We hypothesized that these trajectories represent erroneously tracked trajectories (e.g. movements of two different cells concatenated in error) and reasoned that high displacement kurtosis values in this cluster result from unrealistic single-frame displacements within the trajectory. We chose to eliminate these trajectories from further analysis because they likely represent artifacts in the imaging acquisition. To eliminate these trajectories, we developed a rule to eliminate trajectories with outlier displacements according to the formula below,

$$d > Q3 + t * (IQR) \Rightarrow \text{outlier}$$

where *d* is a displacement in the trajectory, Q3 represents the third quartile of the displacement distribution, *t* is a tunable threshold coefficient, and IQR is the difference between the third and first quartile of the data. We tested values of *t* ranging from 1.5 – 4.5 and plotted the distribution of non-outlier and outlier trajectory displacement kurtoses for each threshold (**Supplementary Figure 2B**). We found that a threshold of 2.5 minimizes the number of eliminated trajectories while also eliminating trajectories with high displacement kurtosis from the dataset. After trajectory elimination, we found that most of the cluster with high displacement kurtosis was eliminated from the data (**Supplementary Figure 2C**). The cleaned dataset consisted of 6,912 trajectories.

After eliminating outliers from the cleaned dataset, we were curious whether shape properties of the displacement distribution were still appropriate measures of these trajectories. In theory, if trajectories do not contain significant outlier displacements, then remaining trajectories with higher displacement kurtosis or skewness values may be an artifact of low sampling frequency. We found that ~2% of trajectories within the cleaned dataset had a displacement skewness > 1.5 and ~2.5% of trajectories had a displacement kurtosis > 2. To test whether these high shape values were an artifact of low sampling frequency, we performed Monte Carlo simulations in which we randomly sampled a normal distribution with the mean and variance of the displacement distribution for each trajectory within our cleaned dataset (**Supplementary Figure 3**). We found that at a sampling frequency of 19 frames, the percentage of trajectories with displacement skewness > 1.5, respectively, were significantly less than 2%, suggesting that these values of displacement skewness are not an artifact of low sampling frequency. On the other hand, the percentage of simulated trajectories at a 19-frame sampling rate with displacement kurtosis > 2 was 1.6% suggesting that a large proportion of remaining experimental trajectories with high displacement kurtosis may result from a low sampling rate in the data. Therefore, we chose to omit kurtosis features from further analysis of the data, which reduced the overall number of motility features being considered by 5 features (36 to 31 total features).

**SUPPLEMENTAL TEXT S2. Quantification of how spatiotemporal resolution impacts simulated motility features.**

We ran single-cell simulations for a fixed MP = 0.95 and TP = 0.0 for several combinations of spatial resolutions (4 μm, 2 μm, 1 μm per voxel dimension) and temporal resolutions (30, 15, 7.5, and 5 seconds per timestep). To ensure accurate comparison with experimental trajectories, we down-sampled simulated trajectories to the framerate of the experimental trajectories before motility feature extraction. For each spatiotemporal resolution, we quantified the fold change in values for displacement and tortuosity features with respect to trajectories generated using a spatial resolution of 4μm per voxel dimension and a temporal resolution of 30 seconds per timestep (**Supplementary Figure 5A**). When cells were moving unidirectionally, the values of displacement features increased proportionally with temporal resolution— i.e., doubling the temporal resolution roughly doubled the average speed or quadrupled the mean squared displacement. This phenomenon arises because increasing the temporal resolution by a factor of 2 (30 to 15 seconds per time step) doubles the required run time to recapitulate a 10-minute experimental trajectory and, accordingly, doubles the possible number of moves a cell agent could make. Therefore, doubling the temporal resolution can double the maximum simulated average speed. Conversely, the values of displacement features decreased proportionally with spatial resolution— i.e., increasing the spatial resolution by a factor of 2 (e.g. from 4μm to 2μm per voxel dimension), halved the distance cell agents travel per timestep and accordingly reduced the average speed by a factor of 2 and the mean squared displacement at 1-minute intervals by a factor of 4. Spatiotemporal resolution appeared to have no effect on simulated trajectory progressivity.

We next repeated this analysis for a fixed MP = 0.95 and TP = 0.9 to quantify how spatiotemporal resolution affects motility features when the cell agent is also turning (**Supplementary Figure 5B**). We found that the magnitude changes in displacement features in response to increasing temporal resolution were less than those compared to when the cells are moving unidirectionally (**Supplementary Figure 5A**). This result likely arises due to down-sampling of simulated trajectories that were generated at higher temporal resolutions than the experiment. Due to this down-sampling, at temporal resolutions greater than the experimental framerate, high TP values could lead to turning behaviors that reduce the quantified displacement of the down-sampled trajectory, but not the trajectory at the simulation framerate. The effect of spatial resolution did not change in response to increasing TP. As before, spatiotemporal resolution appeared to have minimal effect on progressivity values.

Finally, we wanted to assess the role of spatiotemporal resolution globally for several (MP, TP) combinations. For each spatiotemporal resolution pair, we generated a “simulated trajectory space” by aggregating trajectories from simulations run with different MP and TP values between 0 and 1. By globally sampling MP and TP for each spatiotemporal resolution pair, we can assess the entire possible universe of trajectories that each spatiotemporal resolution can generate (**Supplementary Figure 5C-D**). As our prior analyses suggested (**Supplementary Figure 5A**), increasing the spatial resolution of the simulation (i.e. decreasing the size of each voxel dimension) decreased the range of movement of the cell agents. Simultaneously, increasing the temporal resolution (i.e. decreasing the size of the simulation timestep) expanded the range of movement of the cell agent (**Supplementary Figure 5C**). Interestingly, at temporal resolutions higher than the frame rate of experimentally measured trajectories, the average speed of simulated trajectories became dependent on both MP and TP (**Supplementary Figure 5C**). This result arises because when down-sampling simulated trajectories at temporal resolutions greater than the experimental framerate, high TP values could lead to turning behaviors that reduce the overall displacement of the down-sampled trajectory. Therefore, at high temporal resolutions, the average speed at a given MP decreases as TP increases. Tortuosity features, such as progressivity, did not appear to experience changes in MP or TP dependence with respect to spatiotemporal resolution (**Supplementary Figure 5D**).

**SUPPLEMENTAL TEXT S3. Motility parameters give rise to distinct motility features in single-cell versus multi-cell contexts.**

Multi-cell simulations include additional rules to prevent cell-agent overlap (**Supplementary Figure 6A**). To assess the impact of these rules, we performed multi-cell simulations that sampled the same MP and TP parameter space as previously considered in the single-cell simulations, with a fixed spatial resolution of 4μm per voxel dimension and a temporal resolution of 30 seconds per simulation timestep (**Supplementary Figure 6B-E**). As in single-cell simulations, the average speed and MSD at 1-minute intervals increased directly with MP in multi-cell simulations. However, for a given value of MP*,* trajectories from multi-cell simulations appeared to have lower displacement than those from single-cell simulations (**Supplementary Figure 6B-C**). This phenomenon likely arises because in multi-cell simulations, cell agents may fail to move on some timesteps due to obstructions to movement. Therefore, for a given MP value, cell agents in multi-cell simulations effectively make fewer moves than those within single-cell simulations. The degree to which displacement in the multi-cell simulation was less than that in the single-cell simulation was also TP-dependent. For a given (MP, TP) combination, the log fold-change between multi-cell and single-cell values for displacement increased as TP decreased (**Supplementary Figure 6B-C**). This trend likely arises because low TP values correspond to more persistent movement decisions. When a cell agent that moves persistently in the multi-cell simulation faces an obstruction in the lattice space ahead, that agent will move to a lattice space that is a shared neighbor of the current lattice space and the lattice space ahead (**Supplementary Figure 6A**). In this scenario, number of choices to move to a non-obstructed space is fewer than if the cell were to move non-persistently and could turn to any of the surrounding lattice spaces. As a result, the cell agent is even less likely to move at low TP values for the same MP value.

As in single-cell simulations, progressivity values of simulated trajectories in multi-cell simulations decreased as TP increased (**Supplementary Figure 6D**). Multi-cell simulations appeared to generate a lower range of progressivity values (0.23-0.81) than single-cell simulations (0.23-1.0). This trend also likely arises due to the cell agents' decisions to avoid obstructions during persistent movement (i.e., if a cell agent that chooses to move persistently in the multi-cell simulation faces an obstruction in the lattice space ahead, that agent will move to a lattice space that is a shared neighbor of the current lattice space and the lattice space ahead). While this decision minimizes turn angles, this action still results in less-direct movement, which lowers the progressivity value. These decisions to avoid obstructions during persistent movement account for lower observed progressivity values for lower TP values in the multi-cell simulation compared to the single-cell simulation. In both single-cell and multi-cell simulations, the net distance of simulated trajectories increased for higher MP values and decreased for higher TP values (**Supplementary Figure 6E**). The maximum net distance in the multi-cell simulation was 70μm, while the maximum net distance in the single-cell simulation was 103μm. The lower net distance observed in the multi-cell simulation is consistent with the observation that decisions to avoid obstacles lead to lower movement for a given MP value, and even lower movement at lower TP values.

Taken together, these results indicate that cells moving within multicellular environments exhibit fundamentally different motility behaviors for a given (MP, TP combination) to avoid surrounding obstructions. These observations indicated that parameterization for MP and TP should be conducted within multi-cell simulations that reflect how obstructions distort cell motility, rather than in single-cell simulations.

**SUPPLEMENTAL TEXT S4. Comparing mixture modelling with *k*-nearest neighbors.**

In addition to the “top-down”, mixture modelling approach in which we characterized the aggregated distance between each experimental motility behavior and simulated (MP, TP) combination, we assessed a “bottom-up” parameterization approach. In the bottom-up parameterization approach, we instead found the simulated k nearest neighboring trajectories to each experimental trajectory within each motility behavior. We then extracted the distribution of MP and TP that gave rise to these simulated k nearest neighbors (KNN) as the parameter distribution for each motility behavior (**Supplementary Figure 8A**). The motility features distributions for the simulated KNN were highly similar to the experimental ones in each motility behavior (**Supplementary Figure 8B-D**). This high similarity directly results from the simulated KNN being selected from a distance metric that looks for maximum similarity to the experimental trajectory. However, when running multi-cell trajectories that re-sample the (MP, TP) distributions of the simulated KNN this similarity was not reproducible (**Supplementary Figure 8B-D**).

The distribution of displacement features, such as average speed, of the resampled simulated trajectories maintain similarity with each experimental motility behavior (**Supplementary Figure 8B**). However, the distribution of tortuosity features, such as progressivity, no longer accurately fit that of each experimental motility behavior. Specifically, we found that the resampled trajectories under-predicted progressivity values for directed motility behaviors (MB 1, 3, 5; **Supplementary Figure 8C**). The percentage of features across all motility behaviors with a Cohen’s D values > 0.5 increased from ~5% for the simulated KNN to 28% for the trajectories that resampled the (MP, TP) distribution of the simulated KNN. This substantial increase in overall dissimilarity indicates that this bottom-up approach is less robust than the top-down approach. We hypothesized that this lack in robustness arises from the inherent stochasticity in the computational model—each combination of (MP, TP) gives rise to a trajectory with a distribution of displacement and tortuosity values. Therefore, the (MP, TP) value of a simulated neighbor to an experimental trajectory is not guaranteed to reproduce the exact feature values that made that simulated trajectory similar to the experimental one. Indeed, the variability of TP is higher than that of MP because measures of tortuosity are highly sensitive to small deviations in the number of turns the trajectory makes, explaining why the resampled trajectories are particularly dissimilar across tortuosity features and not necessarily displacement features.

We attempted to rectify this issue by selectively re-weighting the (MP, TP) distributions of the simulated KNN toward lower TP values for the directed motility behaviors (see **Methods**). This re-weighting method learned to shift the (MP, TP) distribution for directed behaviors (MB 1, 3, and

5) towards lower TP values. Conversely, this method learned not to reweight the (MP, TP) distribution of tortuous motility behaviors (**Supplementary Figure 8E**). Trajectories that sampled the re-weighted (MP, TP) distributions had slightly improved fits for tortuosity values against the experimental data (**Supplementary Figure 8F-H**). In particular, the median predicted progressivity of these trajectories was more similar to that of the directed motility behaviors (**Supplementary Figure 8G**). The overall percent of features with Cohen’s D > 0.5 across all motility behaviors decreased from 28% to 26%. Therefore, the reweighting approach did not appear to significantly improve model fit against the data. This overall dissimilarity percentage is substantially higher than what we achieved with the top-down mixture modelling approach (14%) presented in the main text.

The mixture modelling approach for optimization (presented in the main text) accounts for the stochasticity of the (MP, TP) distribution by design and yields fits with greater similarity to the experimental data. Thus, we chose to apply this method to derive (MP, TP) combinations for the overall experimental cell types, cell motility classes derived from hierarchical clustering, and cell motility behaviors (derived from *k*-means clustering).

**Supplemental Text S5. Determining run conditions for simulation validation.**

To validate the ability of our simulations to predict GCB-Tfh contact time, we seeded simulations that recapitulated the percentage volume of labelled cells in the simulation to that in each experimental movie. To compute these percentage volumes of GCBs and Tfhs in each experiment, we first examined a spatial histogram of tracked (**Supplementary Figure 9A**) and segmented (**Supplementary Figure 9B**) cell locations across all videos to see whether cells were uniformly dispersed throughout the imaging volume. We found that the imaged cells appeared to only be dispersed across the middle 70% of the imaged *z*-volume. Therefore, we computed the density of tracked and segmented cells as a proportion of seven-ninths of the experimental imaging volume. In the validation simulations, we also included populations of unlabeled GCBs and Tfhs that matched the expected proportion of these cells in the germinal center. The inclusion of these unlabeled cells allows the multi-cell simulations to: 1) account for cell movement based on the abundance of neighboring cell obstructions and 2) predict the expected level of interactions between GCBs and Tfhs within the overall germinal center, which may not be captured due to sparse labeling within the experimental volume.

Because our computational model is stochastic, we next assessed how many repetitions of the model are necessary to achieve reproducible estimates of tracked GCB-Tfh contact time for each experiment. This analysis is particularly important for this set of simulations because the tracked percentage volume of cells is <1% in each experiment, and we are estimating interactions over a very low sample size of cells that is prone to more noise. Simultaneously, to maximize computational efficiency, we tested running simulations at smaller volumes than the experimental volume. At smaller simulated volumes, the noise across simulated repetitions is higher (**Supplementary Figure 9C**). Therefore, reaching a stable estimate in smaller simulated volumes requires more repetitions. Conversely, achieving a stable estimate at larger volumes requires fewer repetitions. For comparison of simulation predictions with experimental measurements, we opted to run simulations at the experimental volume for at least 50 repetitions. For the sake of comparing simulation interaction predictions with experimental measurements, we chose to run simulations at the experimental volume. However, for exploratory analyses, simulations run at smaller volumes with more repetitions appear to reasonably estimate trends at the experimental volume.

**Supplemental Text S6. Predicting motility behavior sampling bias.**

A known limitation of intravital imaging is that by imaging a confined sub-volume of the tissue, cell trajectory measurements are biased towards those that start and end within this imaging volume. We were curious whether the imaging setup biases measurements of more slow-moving trajectories than fast moving trajectories. We hypothesize that the experimental data is more likely to capture slow-moving trajectories that are more likely to stay within the image volume than fast moving trajectories that are more likely to leave the imaging volume (an thus not be tracked for 20 continuous frames). We attempted to apply our simulations to predict the likelihood of each characterized motility behavior leaving the imaging volume. To first validate whether our simulation could do this, we defined a hypothetical bounding box within the imaging volume that was 50% of the imaging volume **(Supplementary Figure 10A)**. We measured the proportion of tracked cells that started within the bounding box that also stayed within the bounding box. We ran corresponding simulations that were the same size as the experimental imaging volume and predicted the proportion of cells that started within the equivalent simulated bounding box and stayed within both the equivalent bounding box and the whole imaging volume. We found that both experimental measurements and simulation predictions suggest that as the speed of the motility behavior decreases the likelihood of staying within the bounded volume increases (**Supplementary Figure 10B**). Moreover, simulation predictions were highly similar to experimental measurements, suggesting that our simulation can predict the likelihood of a given motility behavior staying within a certain volume. We then predicted the proportion of each motility behavior likely to stay within the imaging volume (**Supplementary Figure 10C**). We used these likelihoods to adjust the proportions of simulated motility behaviors for non-tracked cell types in the simulations, representing the phenomenon that non-tracked cells are more likely to capture fast-moving cell behaviors. We found that predictions for labelled GCB-Tfh contact time maintain high correlation with experimental data and were not significantly different from baseline simulations without this adjustment (**Supplementary Figure 10D-E**). Moreover, predictions for all cumulative GCB-Tfh interactions were not changed by this adjustment, suggesting that measurement errors that bias towards slow-moving cells do not significantly impact our measured interactions within this system (**Supplementary Figure 10F-G**).

### SUPPLEMENTARY TABLES.

**Supplementary Table 1. Mouse imaging conditions.** We analyzed 15 time-lapse, intravital images. Each set of images was focused on a single mouse germinal center, with the germinal center light zone comprising the majority of the imaging field. Images were acquired between days 9-11 of the GC reaction. All images contained a single field-of-view. For each set of time-lapse images, we report the imaging day, mouse, and transplanted cell mixture below. Additionally, we report the average number of segmented and tracked cells per image frame within each time-lapse image. These counts were used to estimate labelled cell densities in simulations recapitulating each experiment.

| Image # | Day of GC Reaction | Mouse # | Transplanted cell mixture # | Segmented GCBs | Segmented Tfhs | Tracked GCBs | Tracked Tfhs |
|---|---|---|---|---|---|---|---|
| 1 | 9 | 1 | 1 | 44 | 79 | 8 | 17 |
| 2 | 9 | 2 | 1 | 57 | 97 | 17 | 16 |
| 3 | 9 | 3 | 1 | 112 | 29 | 27 | 2 |
| 4 | 11 | 4 | 1 | 130 | 308 | 32 | 79 |
| 5 | 9 | 5 | 3 | 76 | 67 | 25 | 8 |
| 6 | 10 | 6 | 1 | 41 | 19 | 14 | 3 |
| 7 | 10 | 7 | 3 | 115 | 136 | 37 | 25 |
| 8 | 10 | 8 | 3 | 146 | 90 | 40 | 18 |
| 9 | 10 | 8 | 3 | 135 | 98 | 43 | 23 |
| 10 | 11 | 9 | 3 | 53 | 44 | 17 | 8 |
| 11 | 11 | 10 | 2 | 64 | 69 | 22 | 14 |
| 12 | 11 | 11 | 3 | 77 | 103 | 20 | 17 |
| 13 | 11 | 10 | 2 | 41 | 159 | 14 | 33 |
| 14 | 11 | 13 | 2 | 150 | 108 | 39 | 20 |
| 15 | 11 | 13 | 2 | 113 | 141 | 33 | 26 |
| **Transplanted cell mixture 1:**<br>50% CD4Cre Tomato + 25% Cg1Cre colorless +<br>12.5% Cg1Cre CFP WT + 12.5% Cg1Cre YFP WT | | | | | | | |
| **Transplanted cell mixture 2:**<br>50% CD4Cre Tomato + 25% Cg1Cre colorless +<br>12.5% Cg1Cre CFP WT + 12.5% Cg1Cre YFP Ezh2(Y641F) | | | | | | | |
| **Transplanted cell mixture 3:**<br>50% CD4Cre Tomato + 25% Cg1Cre colorless +<br>12.5% Cg1Cre CFP Ezh2(Y641F) + 12.5% Cg1Cre YFP WT | | | | | | | |

### SUPPLEMENTARY FIGURES

**Supplementary Figure 1. Initial clustering of experimentally-obtained lymphocyte trajectories. A,** Scree plot depicting the cumulative % variance explained by each principal component projection of the data (left). Communality score for each motility feature within a 5-PC representation of the data (right). **B,** Correlation matrix of all the motility features used for clustering of the trajectories. **C,** Two-dimensional UMAP representation of the experimental trajectories with *k*-means cluster labels overlaid. **D,** Projection of standard-scaled average speed (top, left) and non-normalized progressivity (bottom, left) onto UMAP embedding. Corresponding boxplots of average speed (top, right) and progressivity (bottom, right) for each cell type. Statistical testing shown on boxplots indicates results from a Mann-Whitney test. Bar indicates $p<0.05$. **E,** Representative trajectories for each motility behavior (left). Proportion of each motility behavior within each cell type (middle, heatmap) and proportion of each cell type within each motility behavior (right, bar plot). **F,** Projection of standard-scaled displacement kurtosis (top, left) and displacement skewness (bottom, left) onto UMAP embedding. Corresponding boxplots of displacement kurtosis (top, right) and displacement (bottom, right) for each *k*-means cluster. See Supplemental Text S1 for further description of these results.

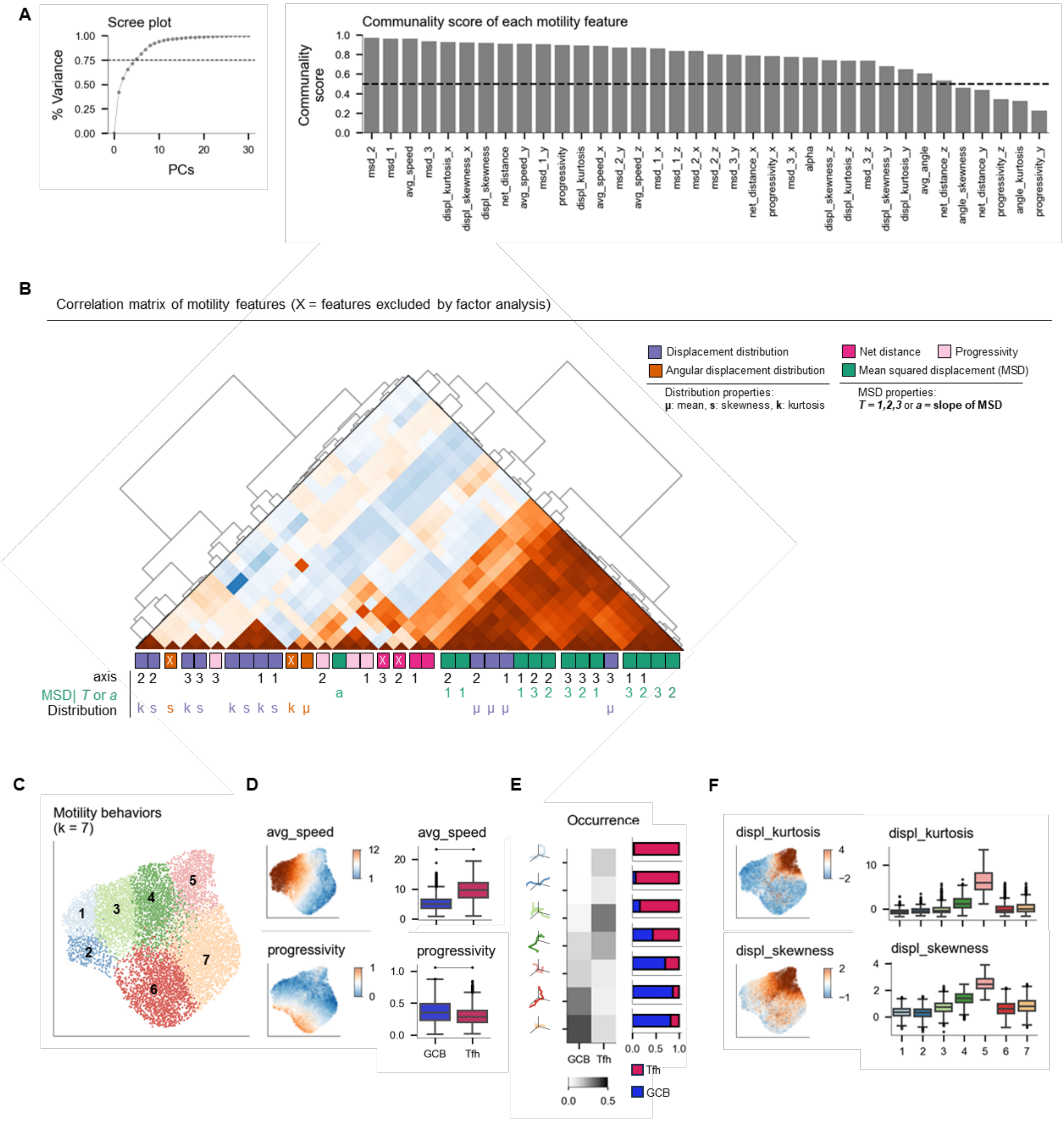


**Supplementary Figure 1. Initial clustering of experimentally-obtained lymphocyte trajectories.**

**Supplementary Figure 2. Examination of trajectories with outlier displacements. A,** Representative histogram of displacements for selected trajectories from initial *k*-means cluster 1, 5, and 6. Cluster 5 tends to have abnormally high displacements that may indicate outliers. **B,** Test of how many trajectories would be discarded for increasing coefficients of the interquartile range term in the outlier decision function. **C,** After eliminating trajectories with displacements greater than the sum of the third quartile and 2.5 times the interquartile range, half of cluster 4 and most of cluster 5, which contained high displacement kurtosis values, were eliminated. See Supplemental Text S1 for further description of these results.

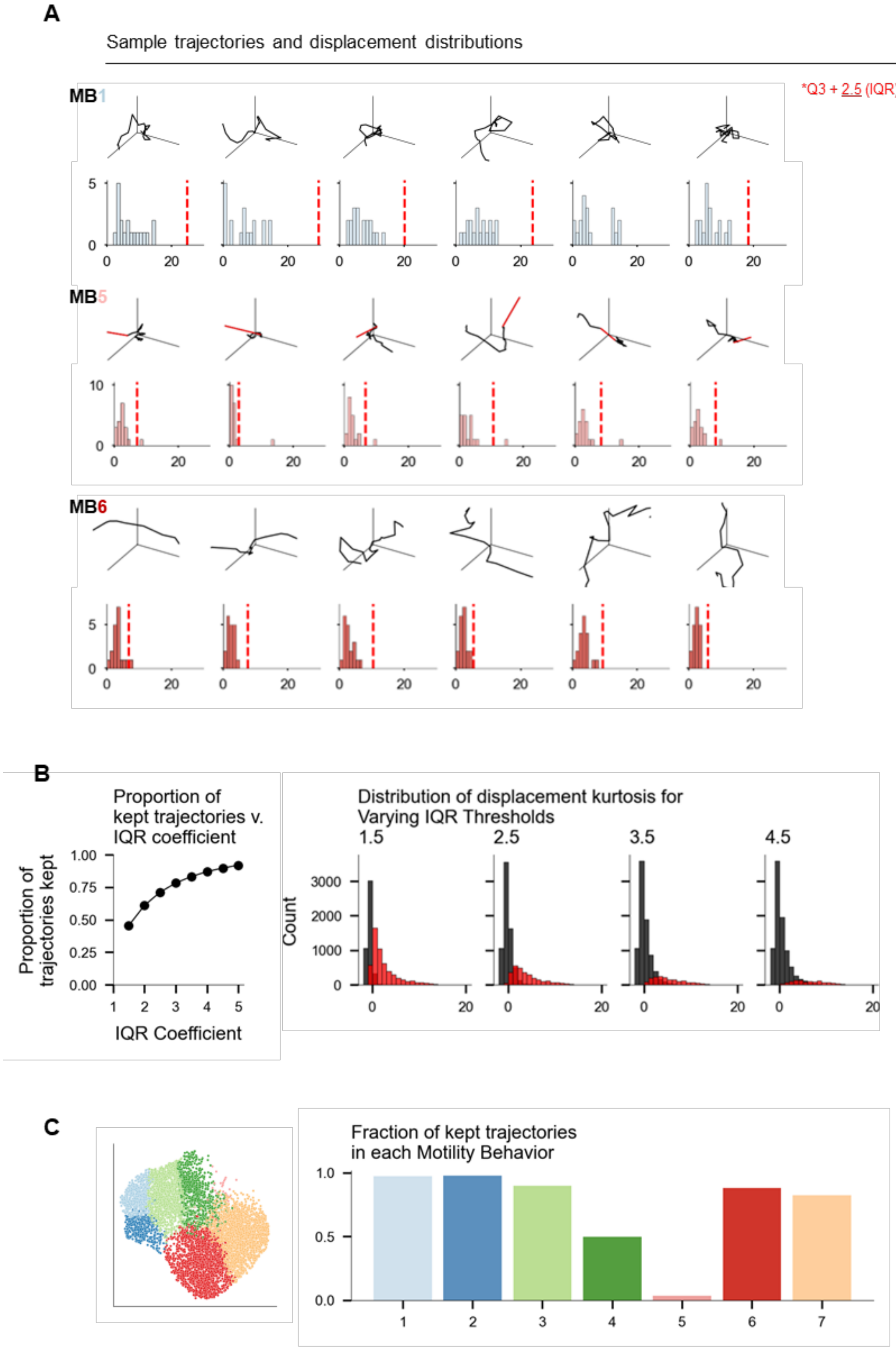


**Supplementary Figure 2. Examination of trajectories with outlier displacements.**

**Supplementary Figure 3. Monte Carlo simulations to predict the likelihood of high displacement skewness and kurtosis values. A,** Distribution of displacement skewness for the experimentally-observed trajectories (left) and trajectories generated from Monte Carlo simulations with increased sampling frequency. **B,** Distribution of displacement kurtosis for the experimentally-observed trajectories (left) and trajectories generated from Monte Carlo simulations with increased sampling frequency. See Supplemental Text S1 for further description of these results.

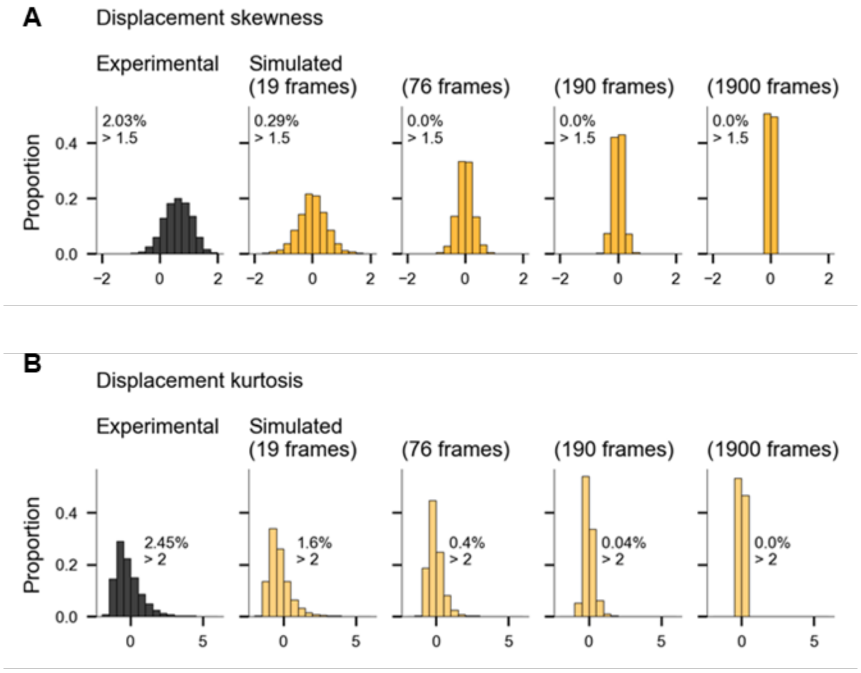


**Supplementary Figure 3. Monte Carlo simulations to predict the likelihood of high displacement skewness and kurtosis values.**

**Supplementary Figure 4. Updated clustering of experimentally-obtained lymphocyte trajectories. A,** Scree plot depicting the cumulative % variance explained by each principal component projection of the data (left). Communality score for each motility feature within a 3-PC representation of the data (right). **B,** Correlation matrix of all the motility features used for clustering of the trajectories. **C,** Elbow plot depicting the average standard deviation of clusters versus the number of cluster (*k*) used for *k*-means clustering. **D,** Concordance plot for chosen 7 clusters. **E,** Radar plot depicting select features for each motility behavior. The clustering described in this figure feeds into Figure 3 in the main manuscript.

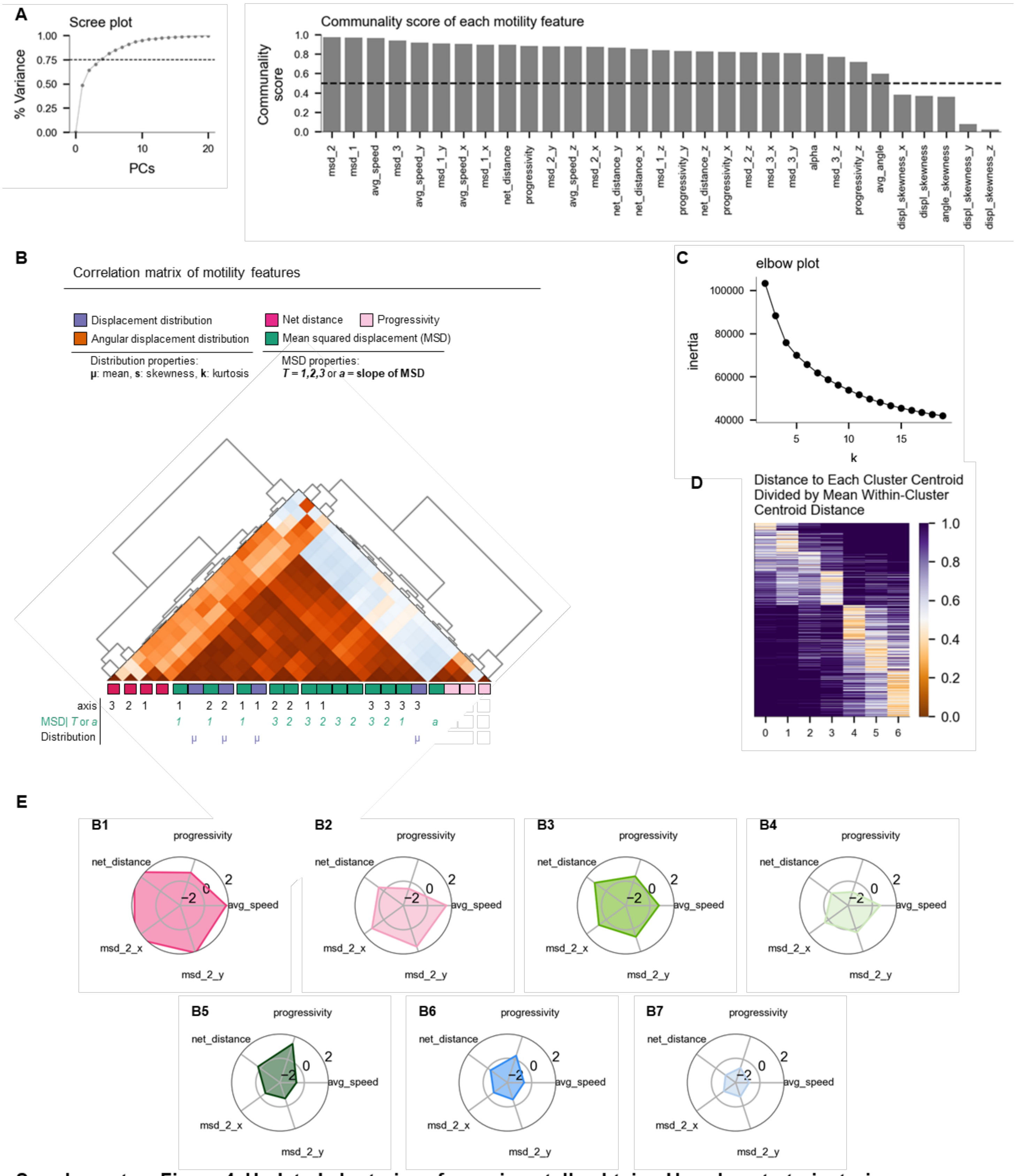


**Supplementary Figure 4. Updated clustering of experimentally-obtained lymphocyte trajectories.**

**Supplementary Figure 5. Global parameter space exploration of motility parameters and spatiotemporal resolution. A-B,** Heatmaps depicting the average $log_2$ fold change between motility features of trajectories generated from varying spatiotemporal resolutions versus a spatiotemporal resolution of 4μm and temporal resolution of 30 seconds per timestep. Analysis was conducted for simulations using MP=0.95, TP=0.0 **(A)** and MP=0.95, TP=0.9 (**B**). **C,** Heatmap showing how average speed varies for varying MP, TP, spatial resolution, and temporal resolution. **D,** Heatmap showing how progressivity varies for varying MP, TP, spatial resolution, and temporal resolution.

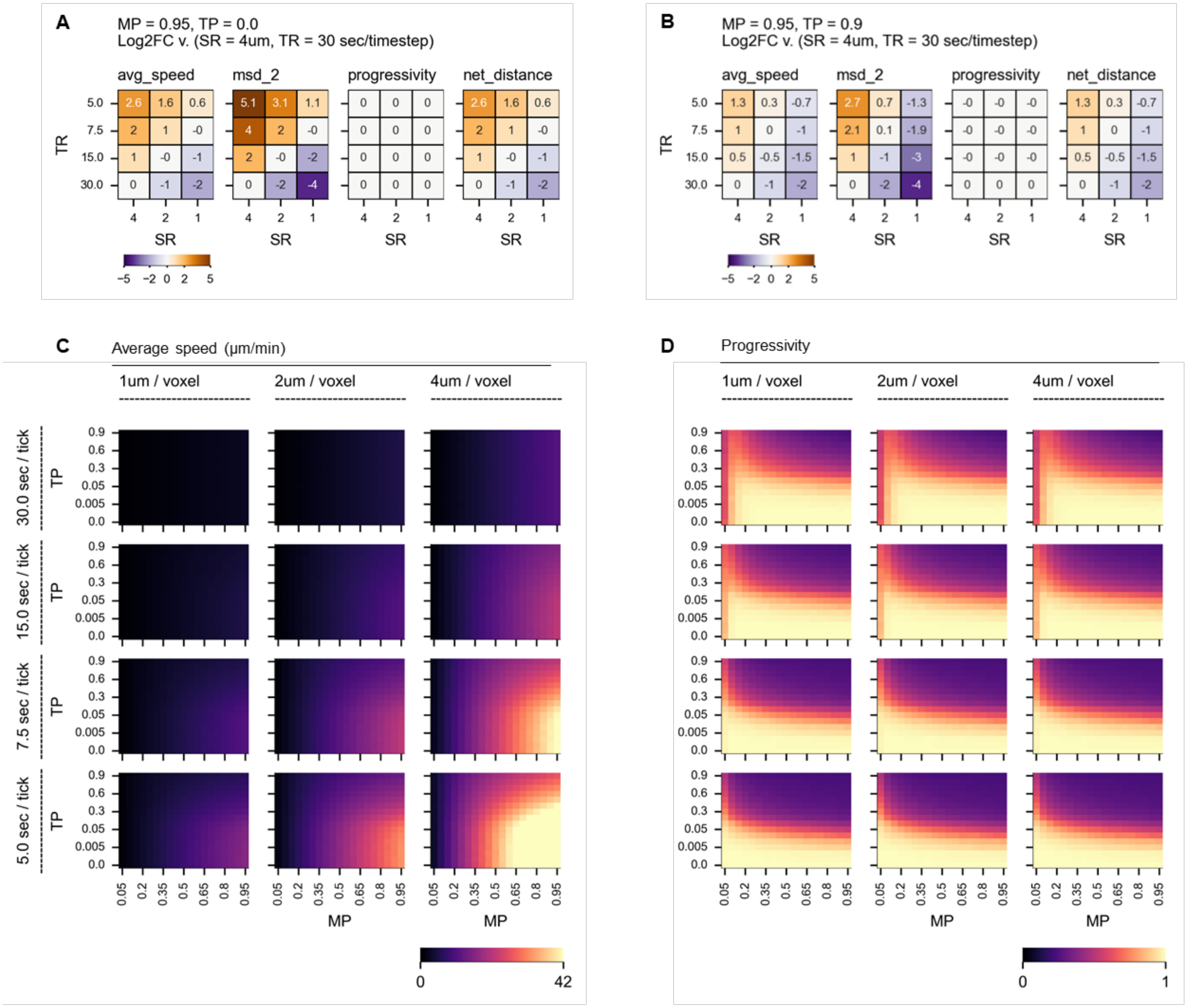


**Supplementary Figure 5. Global parameter space exploration of motility parameters and spatiotemporal resolution.**

**Supplementary Figure 6. Comparison of motility features for a given (MP, TP) combination in single-cell v. multi-cell simulations. A,** Schematic of ABM ruleset for cell-agent movement in single-cell simulations versus multi-cell simulations that require considerations for avoiding obstructions. **B-E,** Heatmaps representing the average value across 2,482 simulated trajectories per (MP, TP) combination in the multi-cell simulation (left) and the $\log_2$ fold change between this average and that for trajectories from the single-cell simulation (right) for the following motility features:  average speed **(B)**, mean squared displacement at 1-minute intervals **(C)**, progressivity **(D)**, and net distance **(E)**.

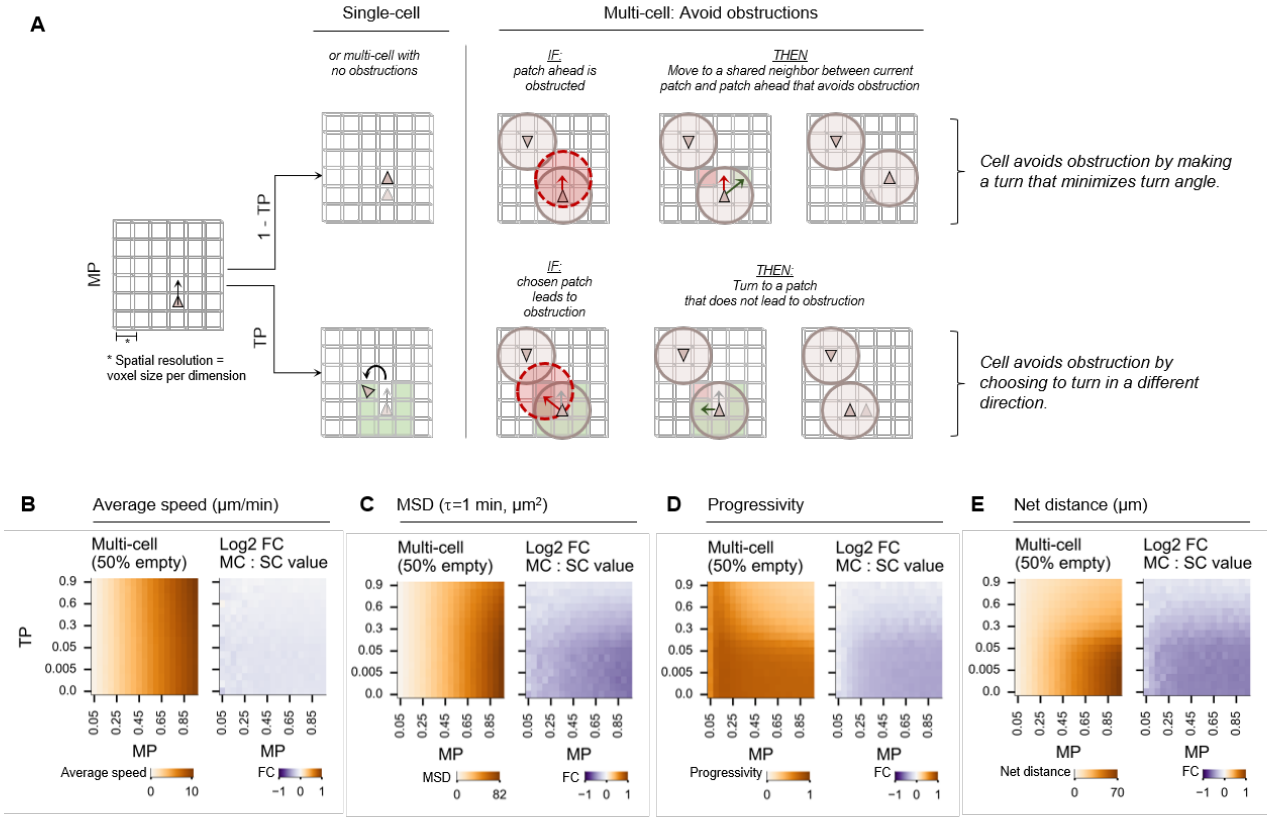


**Supplementary Figure 6. Comparison of motility features for a given (MP, TP) combination in single-cell v. multi-cell simulations.**

**Supplementary Figure 7. Comparison of motility feature distributions between simulated and experimental trajectories for each motility behavior.** Heatmaps depicting the Cohen's D for comparing the simulated versus experimental distribution for each motility feature within each motility behavior. **A-C,** results for parameterization using multi-cell (MC) simulations for the closest (MP, TP) combination (**A**), multi-cell simulations sampling the optimized mixed distribution (**B**), and multi-cell simulations re-sampling the optimized mixed distribution (**C**).

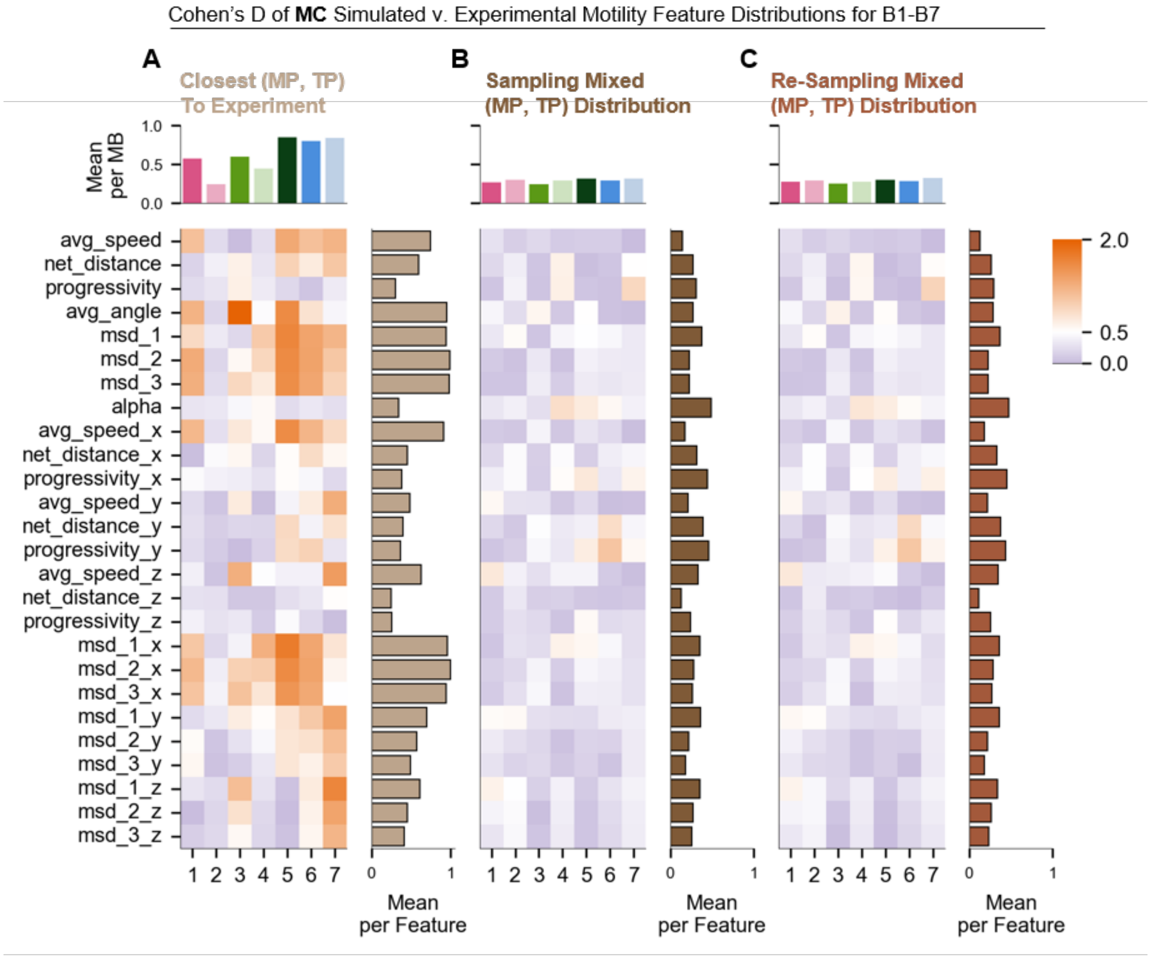


**Supplementary Figure 7. Comparison of motility feature distributions between simulated and experimental trajectories for each motility behavior.**

**Supplementary Figure 8. Bottom-up, k-nearest neighbor parameterization of (MP,TP). A,** (MP, TP) parameter distribution corresponding to *k*=3 simulated neighbors to each experimental trajectory within each motility behavior (MB). **B-C,** Boxplots of average speed (**B**) and progressivity (**C**) distributions of experimentally-observed trajectories, the *k*-nearest simulated trajectories, and simulated trajectories generated by new simulations that re-sample the (MP, TP) parameter distribution derived from the *k*-nearest simulated trajectories. (**D)** Cohen's D for the simulated nearest neighbors to each experimental trajectory and for the trajectories that re-sampled the KNN (MP, TP) distribution. **E,** (MP, TP) parameter distribution derived by re-weighting the original parameter distributions towards lower TP values for more directed motility behaviors (see *Supplemental Text S2*). **F-G,** Boxplots of average speed **(F)** and progressivity (**G**) distributions of experimentally-observed trajectories and trajectories generated by sampling the re-weighted (MP, TP) parameter distributions. **H,** Cohen's D comparing simulated trajectories that sample the re-weighted (MP, TP) distributions with experimentally-observed ones in each motility behavior. See Supplemental Text S2 for further description of these results.

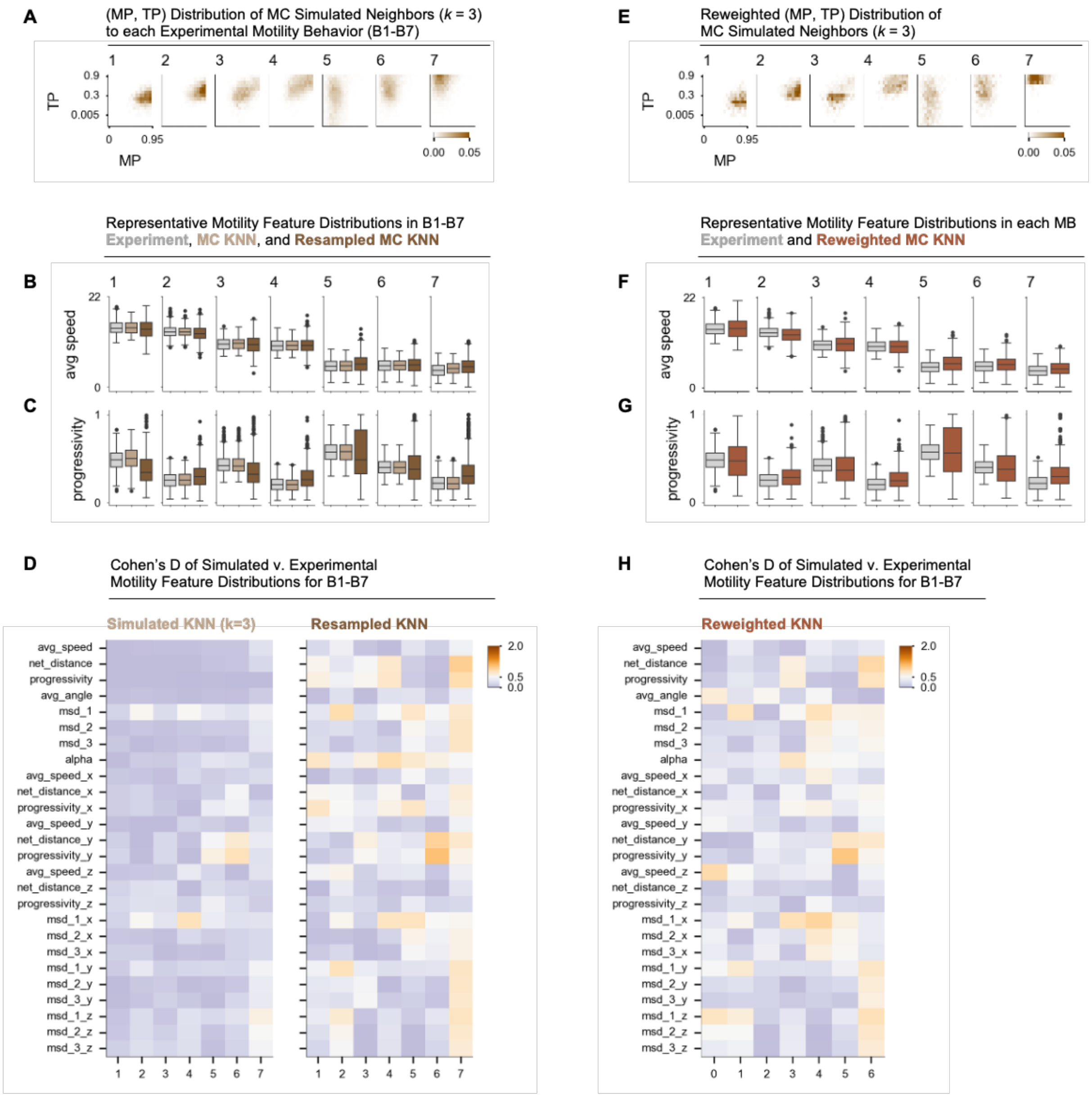


**Supplementary Figure 8. Bottom-up, k-nearest neighbor parameterization of (MP,TP).**

**Supplementary Figure 9. Benchmarking simulation size and repetitions. A,** Heatmap depicting the cumulative count of tracked cells in each 10μm z-frame of the intravital imaging volume. **B,** Heatmap depicting the cumulative count of segmented cells in each 10μm z-frame of the intravital imaging volume. **C,** Cumulative mean of predicted contact time with labelled Tfhs per tracked GCB for simulations run over 100 repetitions at 30%, 50%, and total experimental volumes.

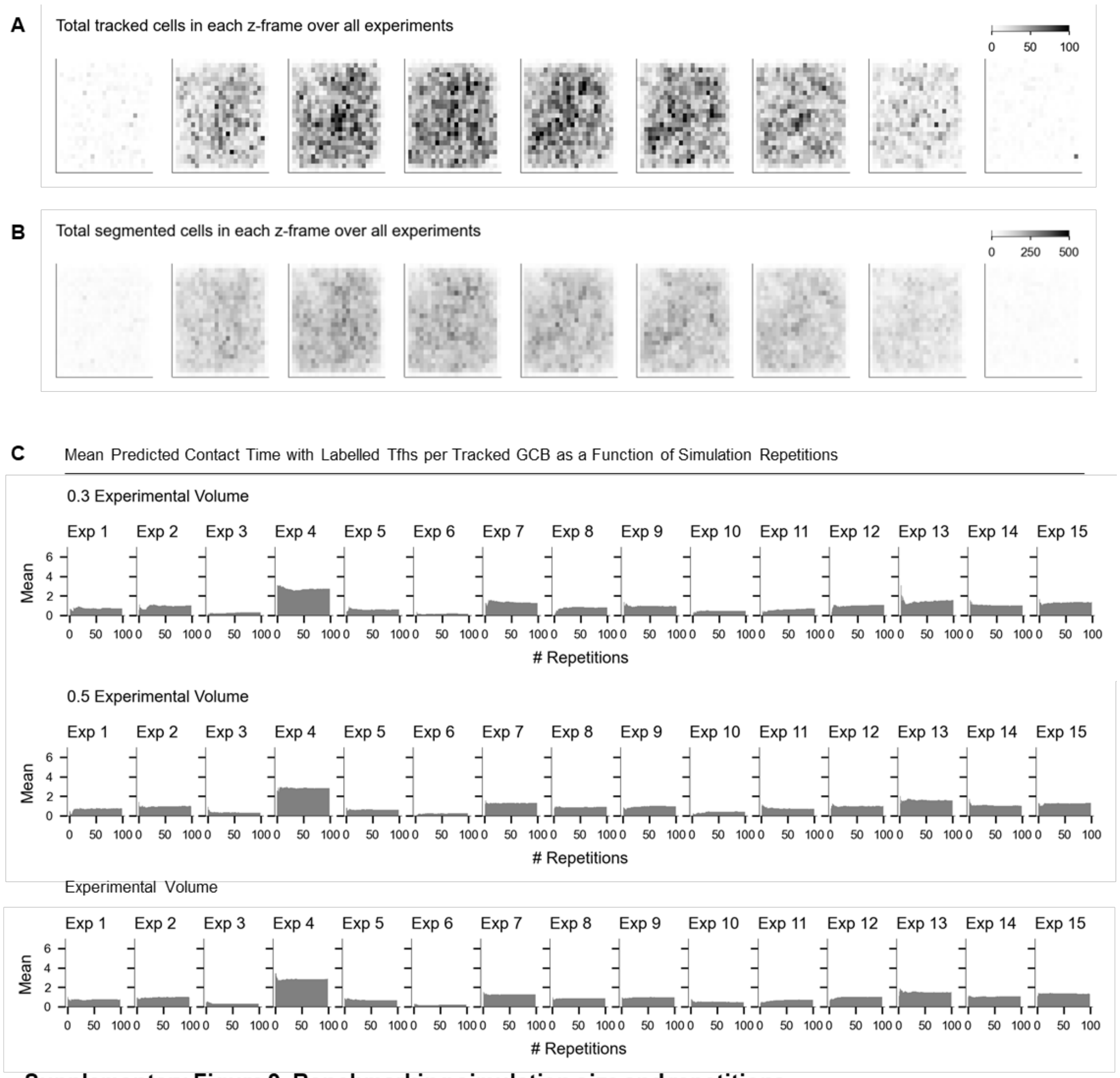


**Supplementary Figure 9. Benchmarking simulation size and repetitions.**

**Supplementary Figure 10. Estimating the likelihood of each motility behavior leaving the imaging volume and assessing impact on predicted GCB-Tfh interactions. A,** Schematic depicting three possible fates for cells in the experiment: tracked cells that stay within the imaging volume and stay within a hypothetical bounding box, tracked cells that stay within the imaging volume but leave the hypothetical bounding box, and untracked cells that leave the imaging volume and are, thus, untracked. **B,** Comparison of experimental measurements and simulation predictions for the proportion of tracked cells that start within a 50% cubic, subvolume centered in the imaging volume that stay within this bounding box. **C,** Predicted proportion of cell agents that stay in the experimental imaging volume for each motility behavior. **D,** Scatter plot of simulation predictions against experimental measurements for contact time with labelled Tfhs per tracked GCB for each motility ruleset. Vertical errorbars represent the standard deviation from 50 simulation repetitions. **E,** Scatter plots of simulation predictions against experimental measurements for contact time with labelled GCBs interactions per tracked Tfh for each motility ruleset. Line indicates a linear regression with the intercept fixed at zero. Simulations were run such that the proportion for each motility behavior was adjusted by the likelihood of leaving the imaging volume **D,** Bar plot representing the $\log_2$ fold change between simulation predictions and experimental measurements for cumulative Tfh contact time per tracked GCB (blue) and cumulative GCB contact time per tracked Tfh (red)**. E-F,** Predicted cumulative Tfh interactions per GCB **(E)** and cumulative GCB interactions per Tfh (**F**) for each motility ruleset. Errorbars represent the standard deviation of 750 simulation repetitions (e.g. 50 per experiment for 15 experiments).Simulations were run at the experimental volume for 50 repetitions per experiment. ‘RMSE’: root mean squared error between average simulation predictions and experimental measurements, ‘r’: r-value for Spearman correlation, ‘p’: *p*-value for Spearman correlation.

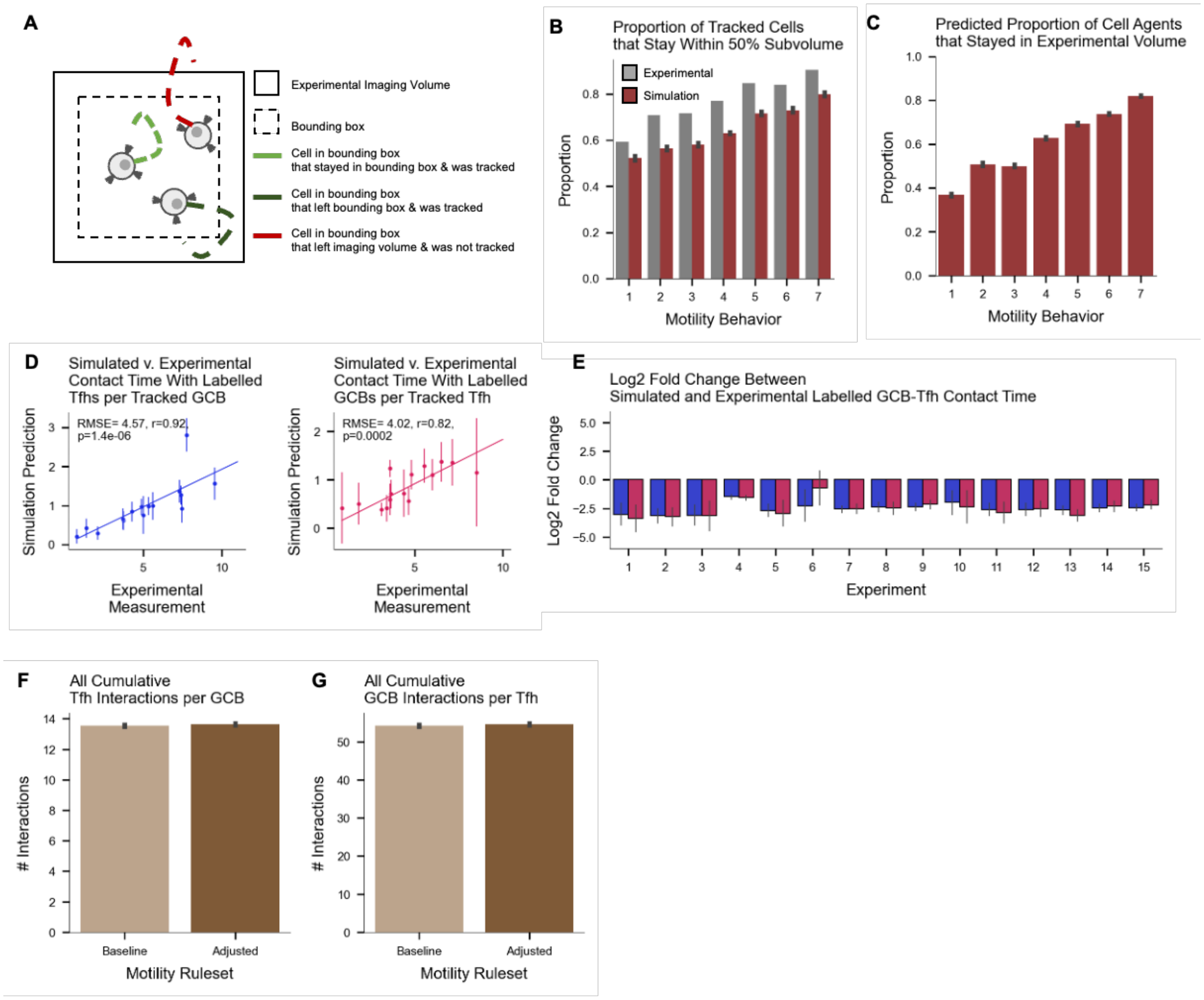


**Supplementary Figure 10. Estimating the likelihood of each motility behavior leaving the imaging volume and assessing impact on predicted GCB-Tfh interactions.**

**Supplementary Figure 11. Simulated cell counts in simulations of static observation volumes with closed boundaries. A,** Total number of simulated GCBs in simulations of a static observation volume for distinct GCB and Tfh motility behaviors. **B,** Total unique Tfh interactions summed across simulated GCBs in simulations of a static observation volume with distinct GCB and Tfh motility behaviors.

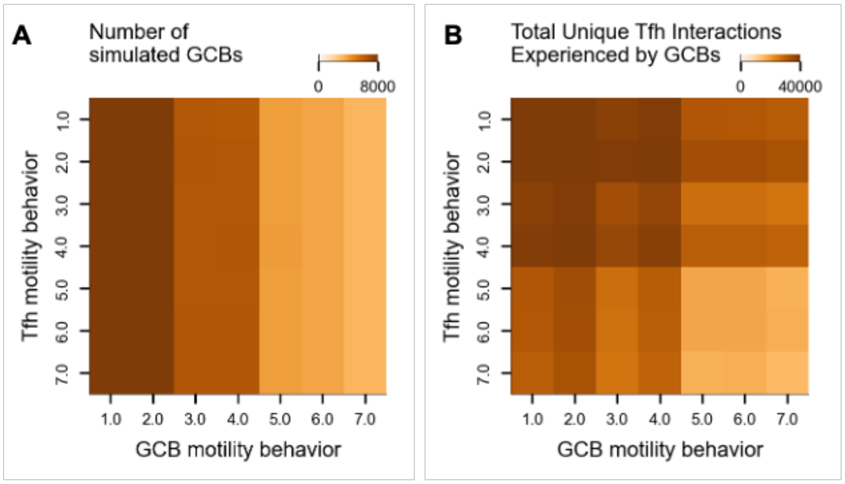


**Supplementary Figure 11. Simulated cell counts in simulations of static observation volumes with closed boundaries.**

**Supplementary Figure 12. Motility embedding showing with wt-EZH2 GCBs and mt-EZH2 GCBs. A,** UMAP embedding from **Figure 3A** with wild-type and mutant GCBs distinguished. **B,** Occurrence of motility behaviors 1-7 across each cell type.

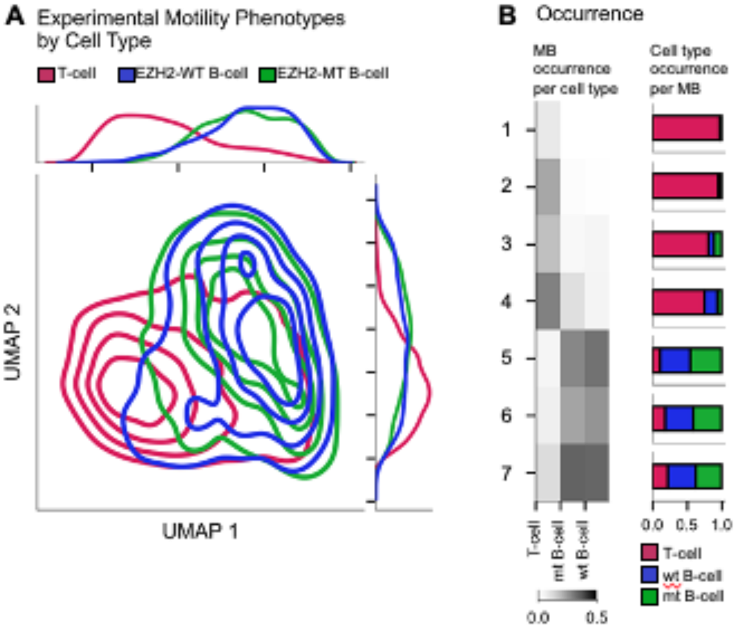


**Supplementary Figure 12. Comparison of wt-EZH2 v. mt-EZH2 GCB motility behaviors.**
